# Enhanced production of heterologous proteins by a synthetic microbial community: Conditions and trade-offs

**DOI:** 10.1101/2020.02.19.955815

**Authors:** Marco Mauri, Jean-Luc Gouzé, Hidde de Jong, Eugenio Cinquemani

## Abstract

Synthetic microbial consortia have been increasingly utilized in biotechnology and experimental evidence shows that suitably engineered consortia can outperform individual species in the synthesis of valuable products. Despite significant achievements, though, a quantitative understanding of the conditions that make this possible, and of the trade-offs due to the concurrent growth of multiple species, is still limited. In this work, we contribute to filling this gap by the investigation of a known prototypical synthetic consortium. A first *E. coli* strain, producing a heterologous protein, is sided by a second *E. coli* strain engineered to scavenge toxic byproducts, thus favoring the growth of the producer at the expense of diverting part of the resources to the growth of the cleaner. The simplicity of the consortium is ideal to perform an in depth-analysis and draw conclusions of more general interest. We develop a coarse-grained mathematical model that quantitatively accounts for literature data from different key growth phenotypes. Based on this, assuming growth in chemostat, we first investigate the conditions enabling stable coexistence of both strains and the effect of the metabolic load due to heterologous protein production. In these conditions, we establish when and to what extent the consortium outperforms the producer alone in terms of productivity. Finally, we show in chemostat as well as in a fed-batch scenario that gain in productivity comes at the price of a reduced yield, reflecting at the level of the consortium resource allocation trade-offs that are well-known for individual species.

## Introduction

Synthetic microbial consortia have been proposed for a variety of applications in industrial and environmental biotechnology and in biomedicine [1–7]. In general, the advantage of synthetic consortia, in comparison with naturally evolved microbial communities, is that they have a simpler composition and often a known interaction structure, which makes them easier to understand and exploit [8]. For example, a co-culture of two engineered *Escherichia coli* strains has been shown to significantly increase the production of muconic acid, a commodity chemical, from a mixture of glucose and acetate [9]. The pathway for the production of muconic acid was distributed over the two strains, coupled by an intermediate metabolite secreted by the first strain and assimilated by the second, which enabled more efficient conversion of the sugar mixture to muconic acid.

Natural communities are a source of inspiration for the design of synthetic consortia. An interesting example is the small ecosystem that emerged during evolution experiments with *E. coli* in a glucose-limited chemostat [10, 11]. In the conditions of the experiment, and more generally when growing fast on glucose, *E. coli* bacteria secrete acetate into the medium [12, 13]. After several hundreds of generations, the initially isogenic population in the bioreactor was found to have differentiated into genetically distinct subpopulations, including a strain with enhanced glucose uptake and acetate secretion rates, and another strain with enhanced acetate uptake and reduced acetate secretion rates [11]. The adaptive genetic changes had reinforced the feeding of one strain on a metabolic by-product of the other, thus enabling their stable coexistence.

Apart from being a model system for studying interactions in naturally occurring microbial communities [14], the results of this evolution experiment also suggest novel applications. The secretion of acetate is not only wasteful but also toxic, as its accumulation in the medium inhibits bacterial growth [15–17]. Growth inhibition by acetate and other weak acids is a well-known problem in industrial biotechnology, where it limits the productivity of strains modified to convert a substrate into a recombinant protein or a metabolite of interest [18, 19]. The *E. coli* community described above suggests an elegant solution to this problem. When coculturing a species producing a heterologous protein of interest and secreting acetate with another species specialized in the assimilation of acetate, the latter could remove the acetate secreted into the medium by the former (Fig. 1). Several instances of synthetic consortia implementing (variants of) this idea have been reported in the literature [20–23]. For example, Zhou *et al.* [23] *describe a consortium of modified E. coli* and *Saccharomyces cerevisiae* strains jointly producing a precursor for an antitumor agent, where the acetate secreted by the bacteria provides the substrate for growth of the yeast population.

**Fig 1.**
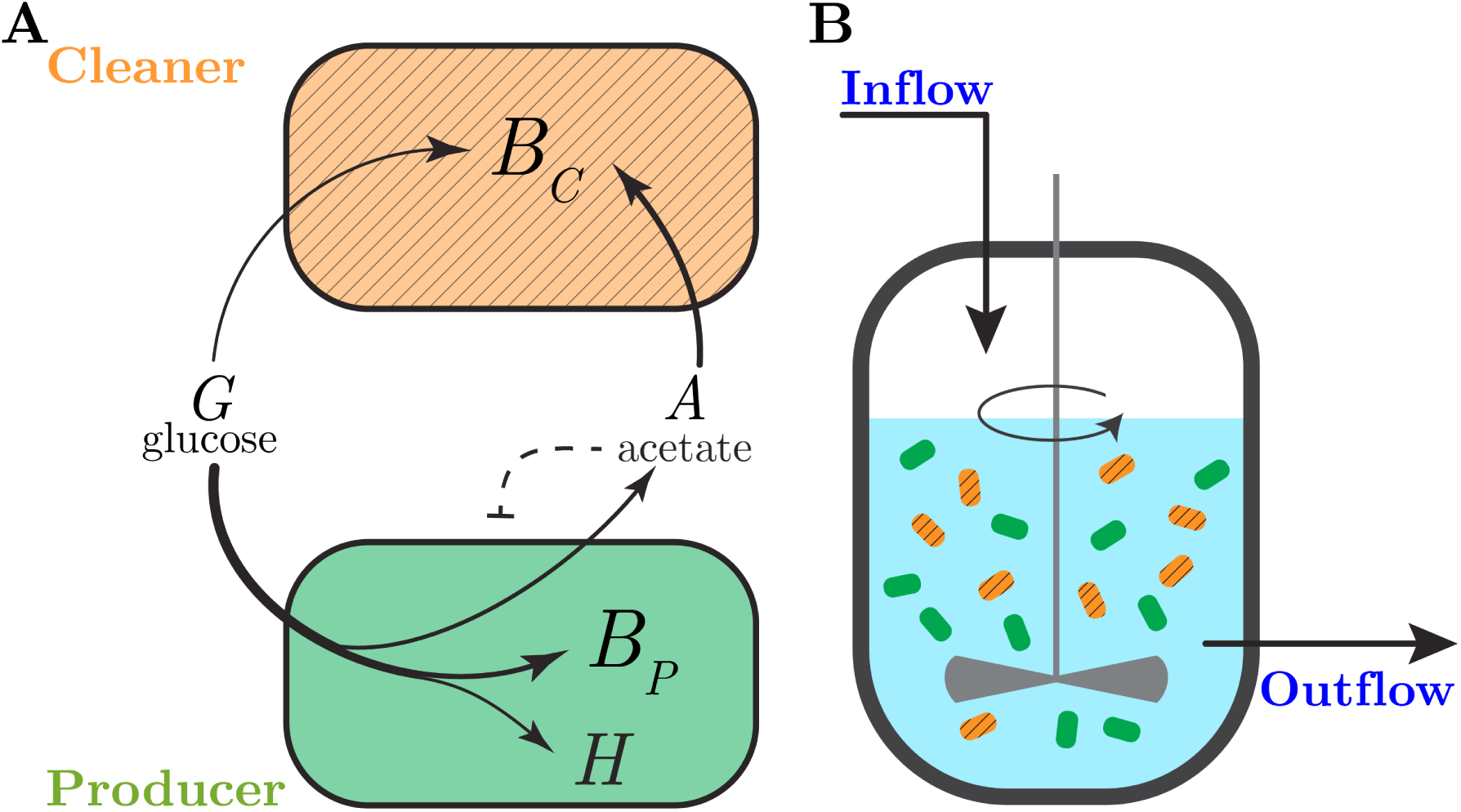
Synthetic consortium for the production of a heterologous protein. (A) The consortium consists of two bacterial species. A first bacterial species growing efficiently on glucose (producer, green) produces a heterologous protein and secretes acetate as a by-product. A second species (cleaner, hatched orange) grows preferentially on acetate, thus removing the latter from the environment and relieving its inhibitory effect on the growth of the producer. Assuming a well-stirred reaction volume, *B*_*P*_ and *B*_*C*_ denote the biomass concentrations of the producer and cleaner species, respectively, *H* is the concentration of the heterologous protein, *A* and *G* denote the concentrations of acetate and glucose. (B) The consortium is grown in a bioreactor, where the inflow and outflow rates and the glucose concentration in the inflowing medium can be externally controlled. We here consider equal and constant inflow and outflow rates, which means that the bioreactor volume is constant and a well-defined steady state can eventually be reached [38].

Despite these promising results, it is not obvious that a consortium of heterologous protein producers and acetate cleaners will produce more of a protein of interest than a single strain. In particular, it requires that the gain in productivity obtained by the depletion of acetate outweighs the loss of substrate incurred by the maintenance of the population of acetate cleaners. The aim of this study is to investigate under which conditions enhanced productivity of a heterologous protein is possible and which trade-offs in the use of available resources this involves. We focus on a prototypical synthetic consortium of *E. coli* strains that was previously implemented and shown to lead to higher biomass accumulation in both batch, chemostat, and biofilm conditions [20].

Mathematical modeling can be leveraged to quantitatively characterize the conditions for improved productivity and inherent trade-offs. For the consortium considered here, and other communities exhibiting cross-feeding and mutualism, a large variety of models have been proposed in the literature [24–27]. We here focus on coarse-grained models that describe by means of a few variables the main processes fueling microbial growth and the interactions between different microbial species. Such models remain mathematically tractable while allowing structural determinants of the productivity of a consortium to be clearly identified.

The models we develop differ in two main respects from existing coarse-grained models of similarly structured communities [28–33]. First, we take into account a range of mechanisms controlling the growth phenotypes of the *E. coli* strains constituting our consortium, with the aim of obtaining quantitatively predictive models. In particular, in addition to acetate toxicity, we consider carbon catabolite repression [34], threshold regulation of acetate overflow [12], and growth-independent maintenance [35]. These mechanisms play an important role in shaping the dynamics of the consortium and are also crucial for making quantitative sense of the experimental data [13]. Second, whereas most of the modeling studies have been concerned with the conditions of coexistence of strains secreting and consuming acetate, we will focus here on the conditions for enhanced productivity of the expression of a heterologous protein by the former. The relationship between coexistence and productivity is not straightforward: while increased productivity requires coexistence, it may also affect the latter, not least through the metabolic load incurred by heterologous protein expression [36, 37].

The models developed in this study will be analyzed in the well-defined steady-state conditions of growth in a chemostat as well as in fed-batch [38]. This results in three main contributions. First, we explain how the overexpression of the heterologous protein modifies the growth rate of the producer strain and thus the boundaries of the producer and cleaner coexistence domain. Second, we determine how productivity is improved under conditions of coexistence, when the loss of substrate due to the growth of the acetate cleaner population is offset by the gain in productivity due to the removal of acetate from the medium and the consequent increase in biomass of the producer. Third, we predict that the increased productivity comes at the price of a reduced yield of conversion of substrate into product, thus generalizing to the community level the rate-yield trade-off observed on the level of individual species [39, 40].

While our study focuses on the synthetic consortium of heterologous protein producers and acetate cleaners described in Fig. 1, the main conclusions are sufficiently general to carry over to other cross-feeding and mutualistic systems harnessed as microbial cell factories. Moreover, the conditions and trade-offs identified here may also provide a starting point for the development of model-based feedback control strategies for optimizing the productivity of synthetic microbial consortia.

## Results

### A dynamical model of the bacterial production strain

#### Model principles and state equations

The synthetic consortium considered here consists of two *E. coli* strains, one growing on glucose and producing the heterologous protein, and the other preferentially growing on acetate and thus removing this growth-inhibitory by-product from the environment [20]. While the overall structure of the model is similar to other population-based models of synthetic mutualistic consortia [28–33], it also differs on key points from previous work in order to account for regulatory mechanisms specific to *E. coli*. We here explain the model for the producer strain in detail and then appropriately modify the model for the cleaner in a later section.

The model for the producer is schematically represented in Fig. 2A. In a well-defined minimal medium, glucose at a concentration *G* [g L^−1^] is taken up at a rate 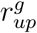 [g gDW^−1^ h^−1^] (where gDW stands for “gram Dry Weight”). The substrate is converted partly into proteins and other macromolecules, which make up most of the biomass, and partly utilized for producing ATP and other energy carriers. In fact, around half of the carbon contents of the substrate is consumed by energy metabolism and leaves the cell in the form of CO_2_ [41]. The conversion of glucose into biomass is thus characterized by the biomass yield coefficient *Y*_*g*_ [gDW g^−1^]. Acetate overflow at a rate 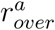 [g gDW^−1^ h^−1^] occurs when the glucose uptake rate reaches a threshold level [42, 43], thus increasing the acetate concentration *A* in the medium. In the absence of glucose, *E. coli* can utilize acetate as an alternative substrate, taken up at a rate 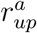 [g gDW^−1^ h^−1^] and converted into biomass with a yield coefficient *Y*_*a*_ [gDW g^−1^]. Accumulation of acetate in the medium inhibits bacterial metabolism and growth [15–17]. The producer strain differs from a wild-type *E. coli* strain in that it carries a plasmid for the inducible expression of a heterologous protein. As a consequence, the total biomass concentration *B*_*tot*_ [gDW L^−1^] is the sum of the concentration of heterologous protein *H* [gDW L^−1^] and the concentration of biomass actively involved in cellular growth and maintenance, the autocatalytic biomass *B* [gDW L^−1^]. The fraction of biomass production assigned to the synthesis of heterologous protein is determined by the (dimensionless) product yield constant *Y*_*h*_. The total biomass *B*_*tot*_ [gDW L^−1^] decays at a rate *k*_*deg*_ *B*_*tot*_, with degradation rate constant *k*_*deg*_ [h^−1^]. In other words, non-growth-related maintenance of the biomass requires an expenditure of resources equal to *k*_*deg*_ *B*_*tot*_ [44].

**Fig 2.**
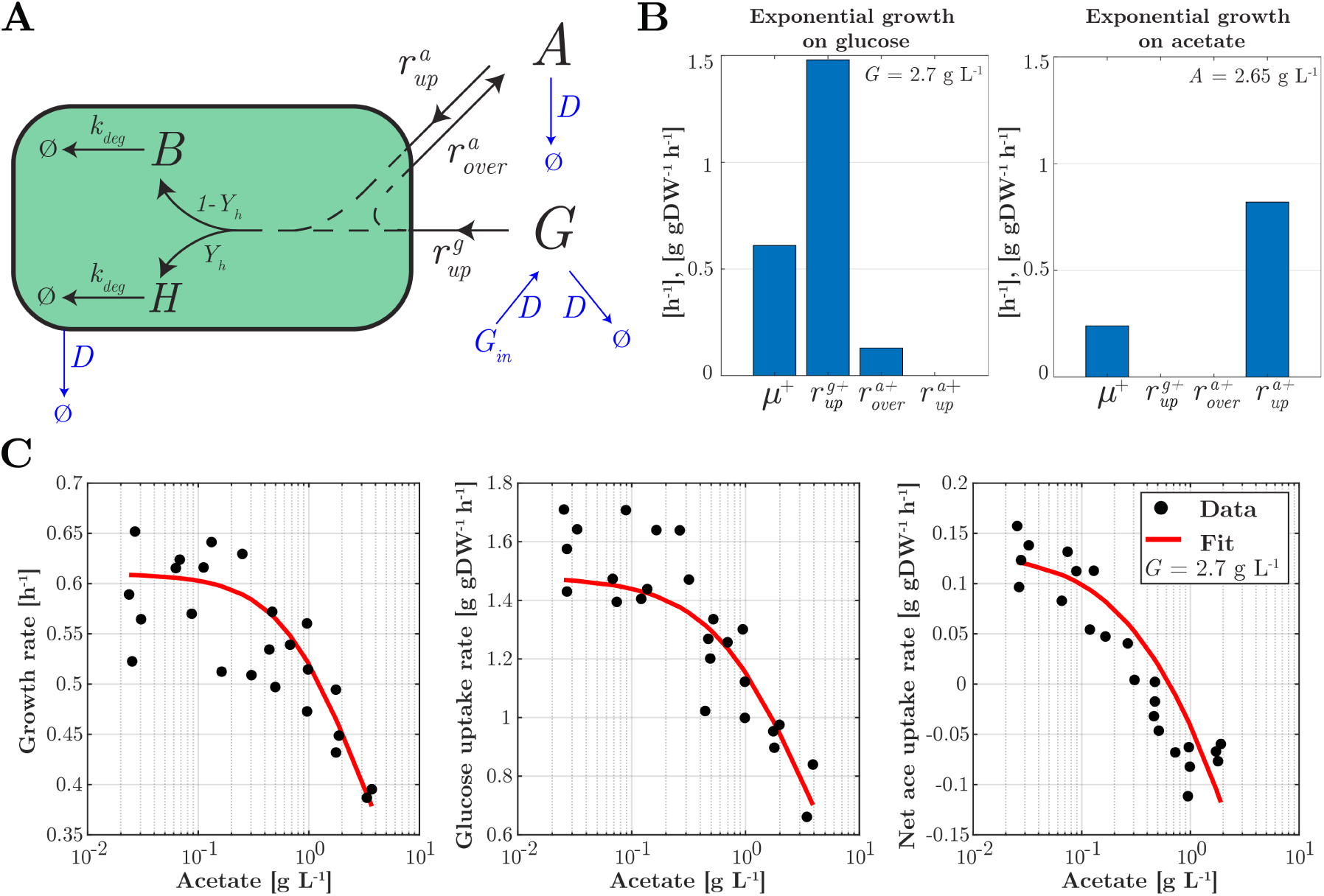
Model of the producer strain and its calibration. (A) The model describes the uptake of nutrients, glucose and acetate, at rates 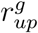 and 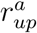, respectively, as well as the pathways for acetate overflow 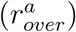, production of autocatalytic biomass and heterologous protein in proportions determined by the yield coefficient *Y*_*h*_, and degradation of biomass (*k*_*deg*_). The bacterial population grows in a bioreactor operating in continuous mode, whereby the dilution rate *D* and the glucose concentration in the inflow *G*_*in*_ can be tuned (indicated in blue). Concentrations in the bioreactor are denoted by *G* for glucose, *A* for acetate, *H* for heterologous protein, and *B* for autocatalytic biomass. For clarity, regulatory interactions due to growth inhibition by acetate and carbon catabolite repression have been omitted from the figure. (B) and (C) Experimental data from [13] used to identify the parameter values of the system of Eqs 1–6. The data include measurements of the growth rate *μ*^+^, the glucose uptake rate 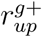, the (net) acetate uptake rate 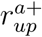, and the acetate overflow rate 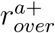 during exponential growth of an *E. coli* wild-type strain in batch conditions in minimal medium with glucose as the sole carbon source (B left panel), acetate as the sole carbon source (B right panel) and in glucose with increasing concentrations of acetate added (C). The + symbol indicates that the measurements have been carried out during exponential growth in batch. The fit of the model to the data is shown as well (red curve). The R^2^ values of the fit are 0.6, 0.77, and 0.77 (from left to right).

The producer strain is assumed to grow in a bioreactor operating in continuous mode [38], that is, with a dilution rate *D* [h^−1^] and a glucose concentration *G*_*in*_ [g L^−1^] in the inflow. This gives rise to the following dynamical system of Ordinary Differential Equations (ODEs):

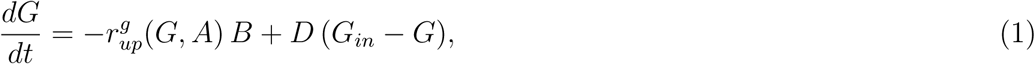

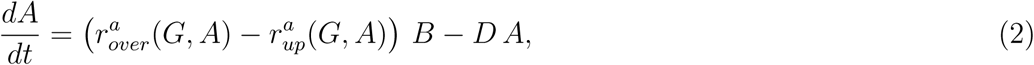

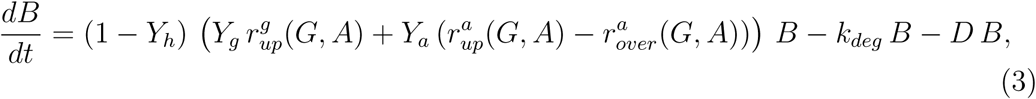

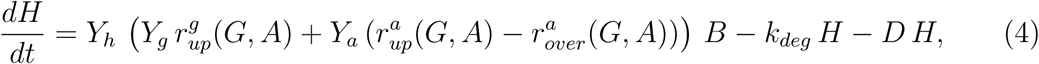

with

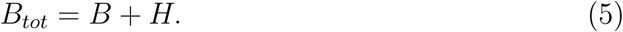

The term 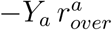 represents the biomass equivalent lost due to acetate overflow, that is, the biomass that would have been produced if the secreted acetate had instead been taken up by the cells. Note that the specific rate of biomass production per unit of biomass, including both heterologous protein and catalytic biomass, is given by 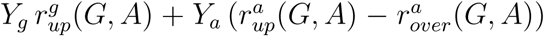. Since only catalytic biomass is actively participating in cellular growth, the total rate of biomass production is obtained by multiplying the specific rate with *B*.

#### Definition of growth rate

The model of Eqs 1-5 allows the (specific) growth rate *µ* [h^−1^] to be explicitly defined in terms of the reaction rates. By definition, the rate of change of the total biomass in the bioreactor equals the biomass produced due to the growth of the bacterial culture (*µ B*_*tot*_) minus the biomass flushed out due to dilution (*D B*_*tot*_):

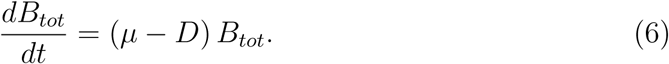

Therefore, the specific growth rate *µ* represents the sum of the rate of change of total biomass per total biomass unit and the dilution rate

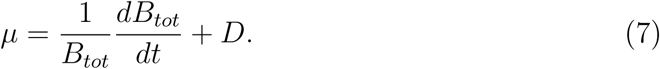

From Eq. 3–5 it then follows that

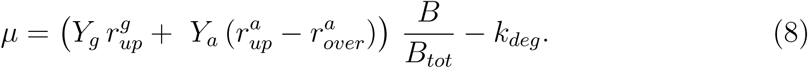

In the absence of heterologous protein production *B*_*tot*_ = *B*, and the growth rate is given by the balance between substrate assimilation and dissimilation and biomass degradation. Heterologous protein production, however, puts a burden on the growth of the population, by drawing away resources from catalytic biomass. Eq. 8 brings out this burden in that a larger *H* leads to a smaller ratio *B/B*_*tot*_ lowering the specific growth rate. At steady state, as shown in the *Methods* section, we find from Eqs 3-4 that 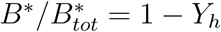, with ^*^ indicating quantities at steady state. Then, Eq. 8 reduces to

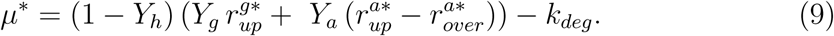

This formulation emphasizes that, at steady state, the growth rate linearly depends on the product yield *Y*_*h*_, as well as on the glucose and acetate uptake rates.

#### Definition of reaction rates

The rate equations are functions of the nutrient concentrations and express the regulatory mechanisms at work. The rate laws are motivated by a combination of modeling assumptions and experimental evidence as explained below.

The uptake rates are defined in such a way that, when inserted into Eq. 8, they lead to Monod-like dependencies of the growth rate on the substrate concentrations, modulated by terms accounting for regulatory effects [45, 46]:

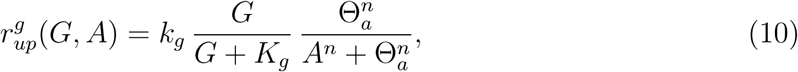

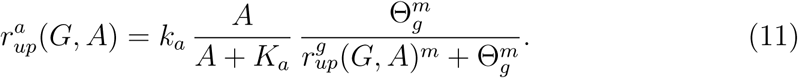

*k*_*g*_, *k*_*a*_ [g gDW^−1^ h^−1^] denote maximal uptake rate constants and *K*_*g*_, *K*_*a*_ [g L^−1^] half-maximal saturation constants. The inhibition term with the constant Θ_*a*_ [g L^−1^] in Eq. 10 represents the inhibitory effect of acetate on bacterial growth [15–17], whereas the inhibition term with the constant Θ_*g*_ [g gDW^−1^ h^−1^] in Eq. 11 corresponds to carbon catabolite repression (CCR), that is, the repression of enzymes in the metabolism of acetate and many other secondary carbon sources while cells are growing on glucose [34, 47]. The strength of CCR is correlated with the phosphorylation state of the preferred glucose uptake system, the phosphotransferase system (PTS) [48], and therefore with the glucose uptake rate in our conditions. Note that, due to acetate inhibition, the glucose uptake rate may be low (and CCR partially relieved) even in the presence of high glucose concentrations in the medium. The exponents *n* and *m* shape the nonlinear effect of acetate inhibition and carbon catabolite repression, respectively.

When the glucose uptake rate exceeds a threshold *l* [g gDW^−1^ h^−1^], *E. coli* produces and secretes acetate as a fermentation by-product [42, 43]. This overflow metabolism has been explained as providing cells, in the presence of excess glucose, with a less efficient but also less costly way to produce ATP than through respiration [43]. Experimental data show that, above the threshold level, the acetate secretion rate is proportional to the excess glucose uptake rate:

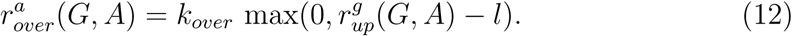

#### Model calibration

The model has 14 parameters most of which have been measured or can be estimated from published data sets. The fact that these experiments have been carried out with different *E. coli* strains and in different conditions carries the risk of obtaining parameter values that are mutually inconsistent. In order to avoid this problem, we have calibrated the model as much as possible against a single, recently published data set for an *E. coli* wild-type strain grown in batch in minimal medium with different concentrations of glucose and acetate [13], that is, for conditions in which *D* = 0 h^−1^ and *Y*_*h*_ = 0. Only the values for the biomass degradation constant *k*_*deg*_, the threshold for acetate overflow *l*, and the half-maximal saturation constants *K*_*g*_ and *K*_*a*_ were taken from the literature. The results of the parameter estimation procedure, described in more detail in the *Methods* section, are plotted in Fig. 2 and show an excellent fit of the model to the data, despite the apparent noise in the measurements, which prevents a perfect fit by any sufficiently smooth model. The parameter values are reported in Table 1 and were compared with other published values for consistency (*Methods*). In order to verify that the resulting model is well-posed, we also performed an *a-posteriori* identifiability analysis (S1 Fig.), showing tight confidence intervals for the parameter estimates. In most simulations, the product yield coefficient was set to *Y*_*h*_ = 0.2.

**Table 1.**
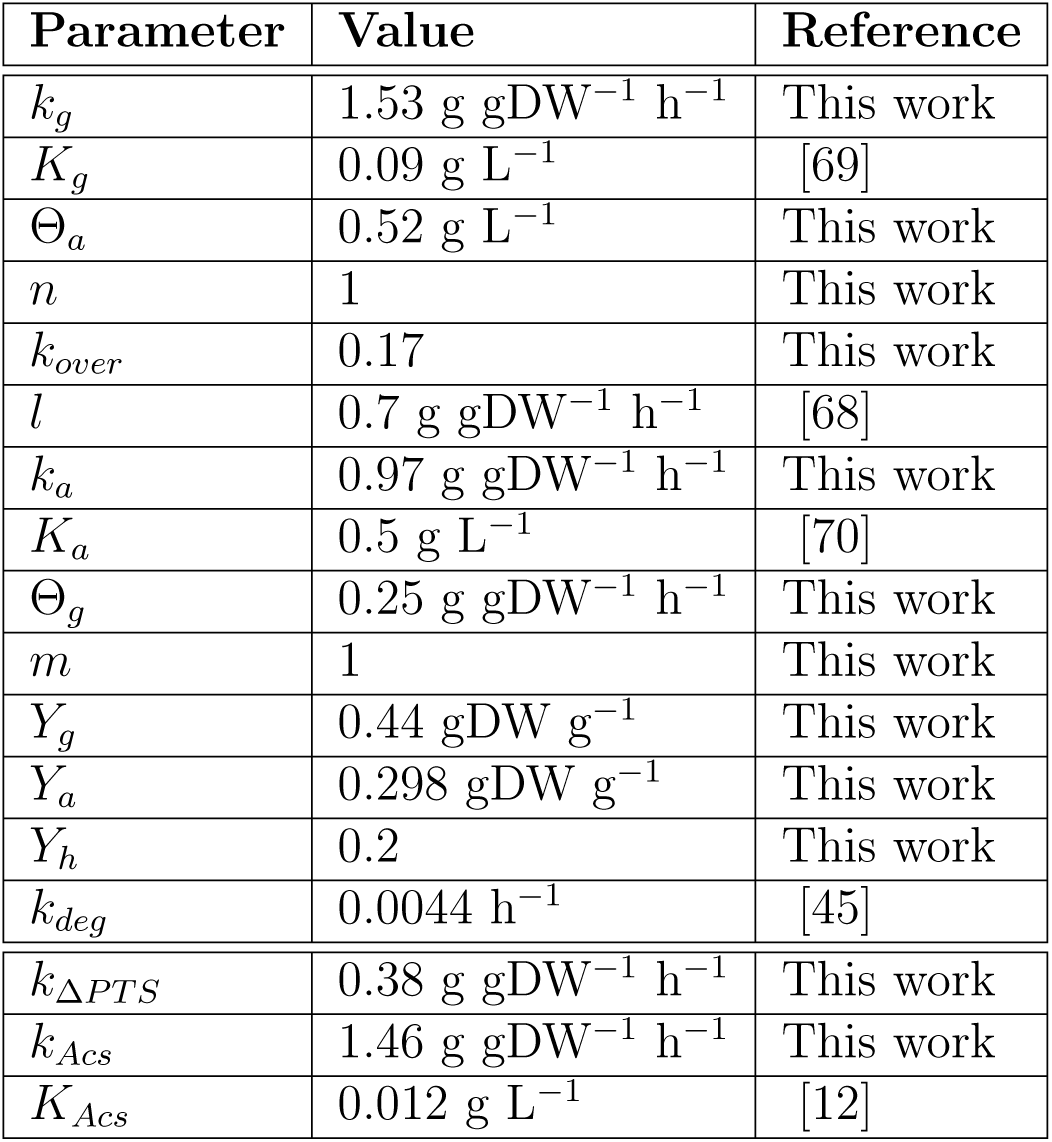
Parameter values. Values of the parameters of the model of Eqs 1–4 describing the producer strain, with rates as in Eqs 10–12, and the cleaner strain, with rates as in Eqs 14–16.

| Parameter | Value | Reference |
| --- | --- | --- |
| $k_g$ | 1.53 g gDW<sup>-1</sup> h<sup>-1</sup> | This work |
| $K_g$ | 0.09 g L<sup>-1</sup> | [69] |
| $\Theta_a$ | 0.52 g L<sup>-1</sup> | This work |
| $n$ | 1 | This work |
| $k_{over}$ | 0.17 | This work |
| $l$ | 0.7 g gDW<sup>-1</sup> h<sup>-1</sup> | [68] |
| $k_a$ | 0.97 g gDW<sup>-1</sup> h<sup>-1</sup> | This work |
| $K_a$ | 0.5 g L<sup>-1</sup> | [70] |
| $\Theta_g$ | 0.25 g gDW<sup>-1</sup> h<sup>-1</sup> | This work |
| $m$ | 1 | This work |
| $Y_g$ | 0.44 gDW g<sup>-1</sup> | This work |
| $Y_a$ | 0.298 gDW g<sup>-1</sup> | This work |
| $Y_h$ | 0.2 | This work |
| $k_{deg}$ | 0.0044 h<sup>-1</sup> | [45] |
| $k_{\Delta PTS}$ | 0.38 g gDW<sup>-1</sup> h<sup>-1</sup> | This work |
| $k_{Acs}$ | 1.46 g gDW<sup>-1</sup> h<sup>-1</sup> | This work |
| $K_{Acs}$ | 0.012 g L<sup>-1</sup> | [12] |

### The model is able to reproduce key growth phenotypes

Simple as it is, the model is able to reproduce a range of experimentally observed growth phenotypes that are important for the study of a consortium aimed at the production of a heterologous protein of interest, both quantitatively and qualitatively. We demonstrate this by validating the model in scenarios that are different from the one used for model calibration (Fig. 2). In particular, we consider dynamic growth in batch of the wild-type strain (diauxic growth on glucose and acetate), exponential growth of an energy-dissipating mutant strain (with a different acetate overflow behavior), and exponential growth of several glucose uptake mutant strains (with different growth rates and acetate secretion rates). In each case, the model predictions are compared with experimental data sets that were not used for identifying the model parameters. Moreover, we explain the relevance of the different scenarios for the construction of the producer and cleaner making up the consortium of Fig. 1.

#### Diauxic growth

When growing in batch in minimal medium with glucose, *E. coli* bacteria attain a high growth rate and secrete acetate into the medium. Only after the glucose has been consumed, the cells start to take up acetate [12,49]. We validated our model by ensuring that it reproduces this so-called diauxic growth behavior by simulating a batch experiment (*D* = 0 h^−1^) in the absence of heterologous protein production (*Y*_*h*_ = 0). The results are shown in Fig. 3A. The biomass concentration in the bioreactor *B* is seen to increase while the glucose concentration (*G*) decreases and acetate accumulates without being consumed (*A*). When the glucose is almost completely depleted (∼ 4 h), cells continue growth on acetate at a lower rate. As shown in the same plot, the simulations are in good agreement with the experimental data of Enjalbert *et al.* [13]. The capability to reproduce diauxic growth depends on the inclusion of a regulatory term for CCR [34, 47] in Eq. 11, a feature absent from many previous models. The property of diauxic growth is preserved in the case of heterologous protein production (S2 Fig.).

**Fig 3.**
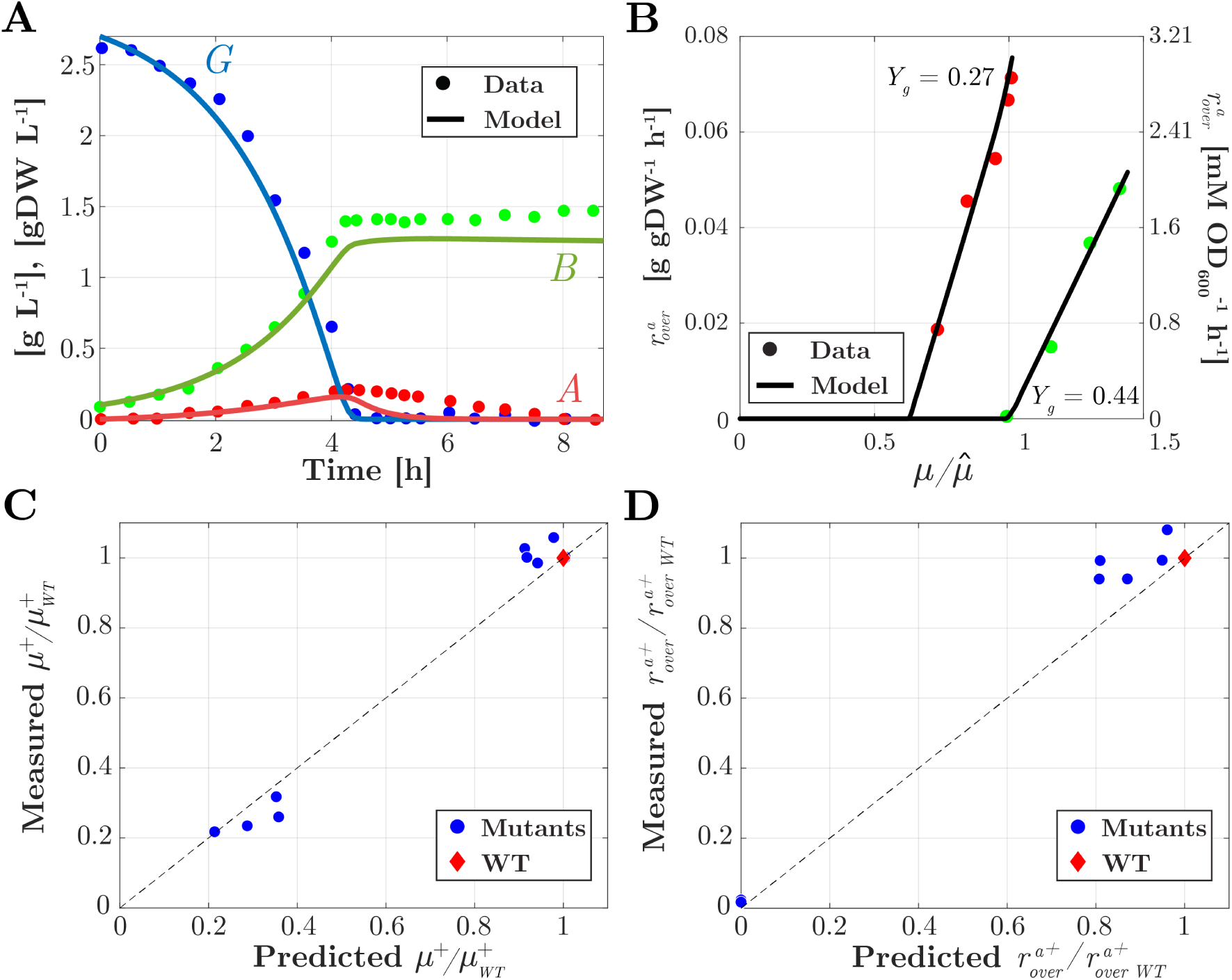
Model validation. (A) Time evolution of the concentration of glucose *G* (blue curve), acetate *A* (red curve), and biomass *B* (green curve) in a bioreactor operating in batch mode, with initial concentrations of 2.7 g L^−1^ of glucose and 0.1 g L^−1^ of biomass. The predicted time-courses are compared with the data from Enjalbert *et al.* [13] (dots). (B) Effect of lower growth yield *Y*_*g*_, accounting for the use of an energy-dissipating LacY mutant, on the onset of acetate overflow, indicated by the overflow rate (black curves) reached at steady state in a bioreactor operating in continuous mode, with *G*_*in*_ = 20 g L^−1^. A decrease in growth yield (*Y*_*g*_ = 0.27 gDW g^−1^ for the mutant vs *Y*_*g*_ = 0.44 gDW g^−1^ for the wild-type strain) causes acetate overflow to occur at a lower growth rate in the mutant and the rate of overflow to depend more strongly on the growth rate (steepness of the curve). The model predictions are compared with the data from Basan *et al.* [43] (dots). Since the onset of acetate overflow occurs at slightly different growth rates in the wild-type strain used for calibration and the wild-type strain used by Basan *et al.*, we compare the relative changes in growth rates upon a decrease in growth yield. 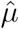 refers to the growth rate at which acetate overflow starts. (C) Predicted versus measured growth rate values for glucose uptake mutants (dots and diamonds, see legend) with different values for *k*_*g*_ that grow exponentially in batch in minimal medium with glucose (G(0)=3.6 g L^−1^). The growth rates of the mutants *µ*^+^ have been normalized with respect to the growth rate of the wild-type strain 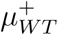 in the same conditions. Dashed lines indicate perfect predictions as a reference. The data are from Steinsiek *et al.* [52]. (D) Idem for acetate secretion rates of mutants 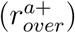 and wild type 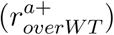. In all four plots there is a good or very good correspondence between the model predictions and the experimental data.

#### Shift in acetate overflow in energy-dissipating mutant

The proton-leaking LacY mutant described by Basan *et al.* [43] has an effect of energy dissipation and therefore a lower growth rate. One might naively expect that the decrease in growth rate would lower the amount of acetate secreted, but this is not the case as energy dissipation causes acetate overflow to start at a lower growth rate [43]. The model reproduces this observation in the case of steady-state growth in a bioreactor operating in continuous mode, without heterologous protein production (*Y*_*h*_ = 0), as shown in Fig. 3B. In order to mimick the effect of the LacY mutation, the growth yield *Y*_*g*_ was decreased from its estimated value of 0.44 gDW g^−1^ (Table 1) to 0.27 gDW g^−1^. Intuitively, the onset of acetate overflow occurs at a lower growth rate, because for a lower growth yield more substrate is needed to produce a given amount of biomass. As a consequence, the glucose uptake rate must be higher to attain the growth rate set by the dilution rate. The model not only correctly predicts that acetate overflow starts at a lower growth rate, but also that the rate of acetate secretion increases more strongly with growth rate, as witnessed by the steeper curve for the lower *Y*_*g*_ value in Fig. 3B. The reproduction of the shift in acetate overflow is a non-trivial achievement of the model and critically depends on the introduction of a regulatory term for acetate overflow in the model, Eq. 12, a feature missing in many other models.

Like energy dissipation, the overexpression of a heterologous protein puts a burden on the metabolism of the host cell, leading to a lower growth rate and other problems for the metabolic engineering of the cell [36, 37]. The analogy between energy dissipation and protein overexpression can be further developed by deriving from Eq. 9 the following relation between the yields *Y*_*g*_, *Y*_*h*_, the dilution rate *D*, and the threshold *l* for acetate overflow:

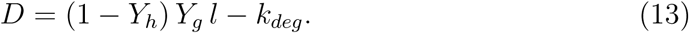

In the derivation we used that *µ*^*^ = *D* and overflow starts at 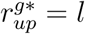, where 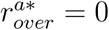 and 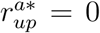 (due to carbon catabolite repression). Eq. 13 shows that a glucose uptake rate equal to *l* occurs at lower *D* for both lower *Y*_*g*_ (energy dissipation) and higher *Y*_*h*_ (heterologous protein production). For an increase in product yield, the model indeed predicts the same shift in acetate overflow as for a decrease in growth yield (S2 Fig.), consistent with reports in the literature [50].

#### Changes in growth rate and acetate secretion rate in glucose uptake mutants

While the PTS is the main glucose uptake system of *E. coli*, a number of other systems are capable of transporting glucose into the cell, such as maltose and galactose transporters [51, 52]. Steinsiek *et al.* systematically tested during exponential growth in batch how the growth rate, the glucose uptake rate, and acetate secretion rate change in strains in which (combinations of) uptake systems have been deleted. The mutants are straightforward to simulate by means of the model bearing in mind that the deletion of an uptake system decreases the value of *k*_*g*_, the maximum glucose uptake rate. More precisely, the observed relative decrease of the glucose uptake rate in a mutant strain leads to a decrease in the same proportion of *k*_*g*_. The predicted growth rates and acetate secretion rates for the mutants correspond well to those measured experimentally by Steinsiek *et al.* (Fig. 3C-D). As expected, for mutants strongly reducing glucose uptake the growth rates are low and no acetate overflow occurs. Mutations in glucose uptake systems form the basis for constructing a cleaner strain that does not or only weakly grow on glucose but instead takes up acetate at a high rate (see [20] and below).

### A dynamical model of the producer-cleaner consortium

As shown in Fig. 3, even at low concentrations, the acetate secreted by the *E. coli* producer strain impairs growth, and therefore the capability to produce the heterologous protein in high amounts. The detrimental effect of acetate could be alleviated by engineering a second *E. coli* strain so as to preferentially take up acetate, the so-called cleaner strain. Below we briefly describe the genetic modifications required to turn a wild-type *E. coli* strain into a cleaner strain, and we provide the model of the synthetic consortium consisting of producers and cleaners. A very similar consortium has been implemented in previous experimental work [20] and coexistence properties have been analyzed by means of a mathematical model of the consortium [31]. The analysis in the next section will focus on a different topic, the conditions and trade-offs underlying improved productivity of a heterologous protein.

#### Design of cleaner strain and model equations

The cleaner strain is obtained from a wild-type *E. coli* strain by knocking-out the gene *ptsG*, encoding a major subunit of the glucose uptake system PTS [51], and transforming this strain with a plasmid enabling the inducible overexpression of the native gene *acs*, coding for the enzyme acetyl-CoA synthetase (Acs) [12]. It has been shown that deletion of *ptsG* reduces the glucose uptake rate 4-fold [52]. The possibility of the cell to continue to take up glucose at a reduced rate, through other, non-specific systems, is advantageous for our purpose, since it allows the cleaner to grow faster than on acetate alone and thus enlarges the coexistence range in continuous-mode cultivation (see below). Overexpressing *acs* has been shown to increase the growth rate of *E. coli* on acetate [53]. Unlike the producer, the cleaner strain does not express the heterologous protein of interest.

With the above design, the model of the cleaner strain is similar to that of the producer, except that Eq. 4 can be eliminated and *B*_*tot*_ = *B*. The rate equations for the cleaner are the following, using the subscript _*C*_ to refer to concentrations and rates of the cleaner:

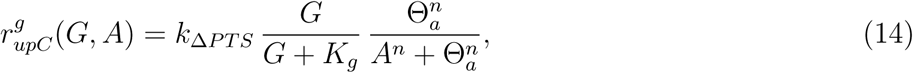

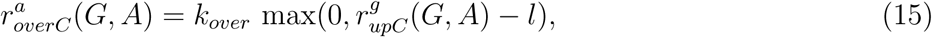

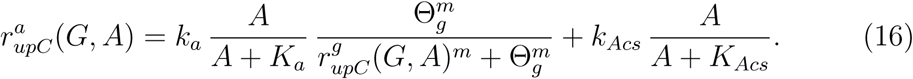

The only modifications with respect to the producer strain are the replacement of *k*_*g*_ by *k*_Δ*PTS*_ [g gDW^−1^ h^−1^], *k*_Δ*PTS*_ < *k*_*g*_, in Eq. 14, and the addition of an extra term acounting for Acs overexpression in Eq. 16, with maximal uptake rate *k*_*Acs*_ [g gDW^−1^ h^−1^], *k*_*Acs*_ > *k*_*a*_, and half-maximal saturation constant *K*_*Acs*_ [g L^−1^].

#### Model of consortium

Next, we assemble the model for the protein producer and the acetate cleaner to obtain the model of the consortium (Fig. 4). The two populations, growing in the same reactor environment, interact in a number of direct and indirect ways. Both the producer and the cleaner take up glucose from the environment, while in addition the cleaner is capable of assimilating acetate secreted by the producer. The removal of acetate cleans the environment from a toxic by-product inhibiting the growth of, especially, the producer. The mutualistic interaction structure of the consortium, in which both strains favor growth of the other, causes the consortium to escape the exclusion principle [54] and makes coexistence of the two strains possible.

**Fig 4.**
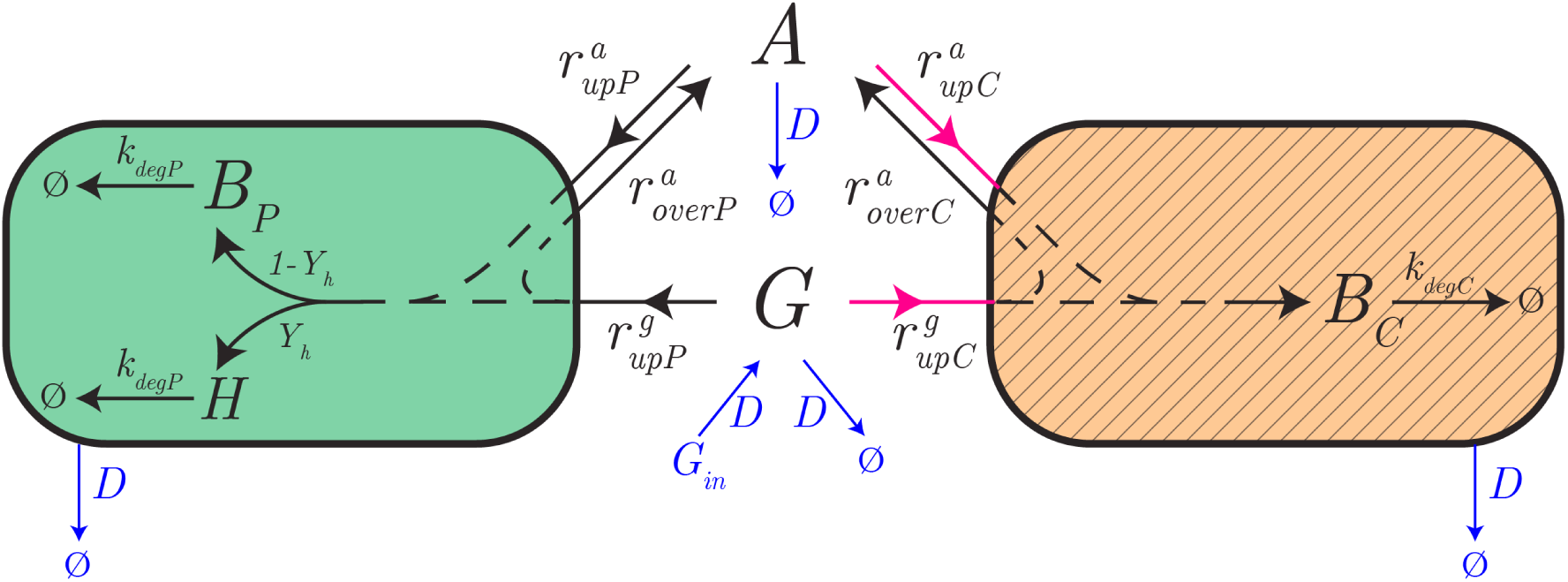
Model of the producer-cleaner consortium. The model describes the protein producer and acetate cleaner strains, in green and hatched orange, respectively. The producer strain preferentially grows on glucose, whereas the cleaner has been modified in such a way as to prefer acetate over glucose. The genetic modifications (purple uptake arrows) include the deletion of the preferred glucose uptake system and the overexpression of a selected enzyme in acetate metabolism. The producer also expresses the heterologous protein of interest. The biomass concentrations of the producer and the cleaner are denoted by *B*_*P*_ and *B*_*C*_, respectively. Reaction rates specific to the producer and the cleaner are also identified by the subscripts _*P*_ and _*C*_, respectively (*e.g.*, 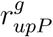 *vs* 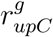 for the glucose uptake rate). The consortium grows in a bioreactor operating in continuous mode, whereby the dilution rate *D* and the glucose concentration in the inflow *G*_*in*_ can be tuned (indicated in thin blue). For clarity, regulatory interactions due to growth inhibition by acetate and carbon catabolite repression have been omitted from the figure.

The dynamics of the system are thus characterized by the following system of ODEs, where the dependency of the rate expressions on the concentrations has been dropped for clarity:

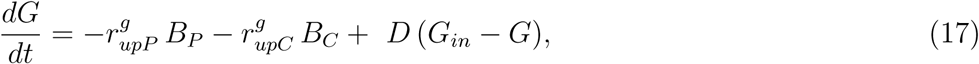

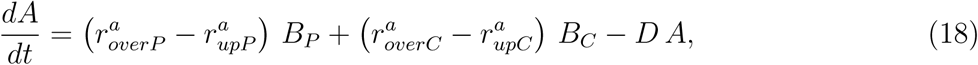

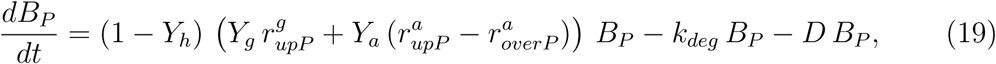

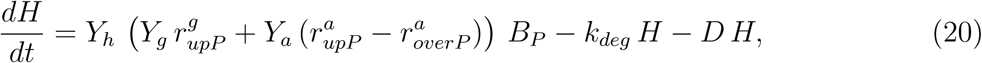

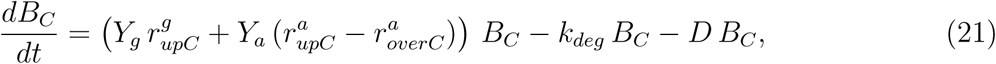

completed by the following mass conservation equation:

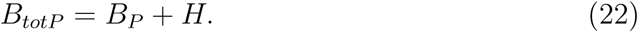

The growth rates of the producer and cleaner strains are defined by the following equations

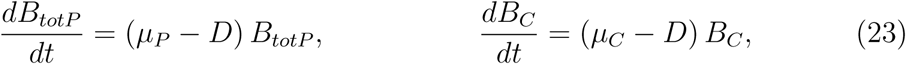

which leads to expressions for *µ*_*P*_, and *µ*_*C*_ analogously to Eq. 8. S3 Fig. shows an example simulation of the consortium, in conditions leading to coexistence of producer and cleaner strains.

### Coexistence of producer and cleaner depends on the product yield

When can the species composing the consortium stably coexist? To answer this question, we study the consortium model developed above in chemostat, that is, we investigate the existence and nature of the (stable) steady states of the model. First of all, setting both derivatives in Eq. 23 to zero, one finds that nonzero steady-state concentrations 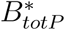 and 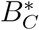 can be obtained only if 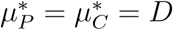. This simple but crucial fact immediately implies that *D* must not exceed the maximal growth rate of either of the species. At the same time, acetate, the primary substrate for cleaner growth, is only available upon overflow in the producer. Since the latter occurs when 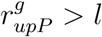, and 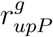 increases with the producer growth rate (see Eqs 9), it follows that persistence of the cleaner is guaranteed only by fast enough growth of the producer. All this suggests that coexistence is only possible in a finite range of growth rates (or equivalently, dilution rates). Since the producer growth rate is affected by the product yield *Y*_*h*_, this range must depend on *Y*_*h*_ as well. In order to quantitatively pinpoint the effects of all these factors on the stability of the consortium, in the following paragraphs we perform steady-state analysis of Eq. 17–21 in a variety of experimental conditions.

#### Dependence of coexistence on *D* and *g*_*in*_

For ease of illustration, in this section, we consider the system with *Y*_*h*_ = 0, that is, in absence of heterologous protein synthesis. All other parameters of the consortium are fixed as in Table 1. The case *Y*_*h*_ > 0 will be considered below.

We sought the (real, nonnegative, stable) steady states of the system for a grid of values of the bioreactor inflow parameters in the region (*D, G*_*in*_) ∈ [0, 0.7] h^−1^ × [0, 20] g L^−1^. Assuming that both biomasses are initially present, we first used a numerical integration approach (see details in *Methods*). For every value of the pair (*D, G*_*in*_), unique steady-state biomass concentrations 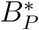 and 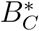 were found for any initial positive biomass concentrations. Uniqueness of the steady state was reconfirmed by an algebraic approach (see *Methods*). Absence of multistability (in particular, bistability) differs from the results in [31] and follows from the monotonic dependence of uptake rates, and therefore growth rates, on substrate concentrations (see *Discussion*).

The results from the numerical computation of 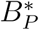 and 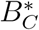 are summarized in Fig. 5A (the same results from the algebraic approach are shown in S4 Fig.). As expected, stable coexistence (nonzero 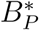 and 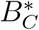) is only observed in a finite range of values of *D*. This range depends very mildly on the specific value of *G*_*in*_ above a small threshold of about 2 g L^−1^. For a given value of *G*_*in*_ above this threshold, four different regimes are encountered through increasing values of *D*, corresponding to stable existence of the sole producer, coexistence, cleaner washout, and producer and cleaner washout. To discuss what characterizes these regimes, we rely on Fig. 5C and 5E, showing steady-state values of several rates and biomass concentrations as a function of *D*.

**Fig 5.**
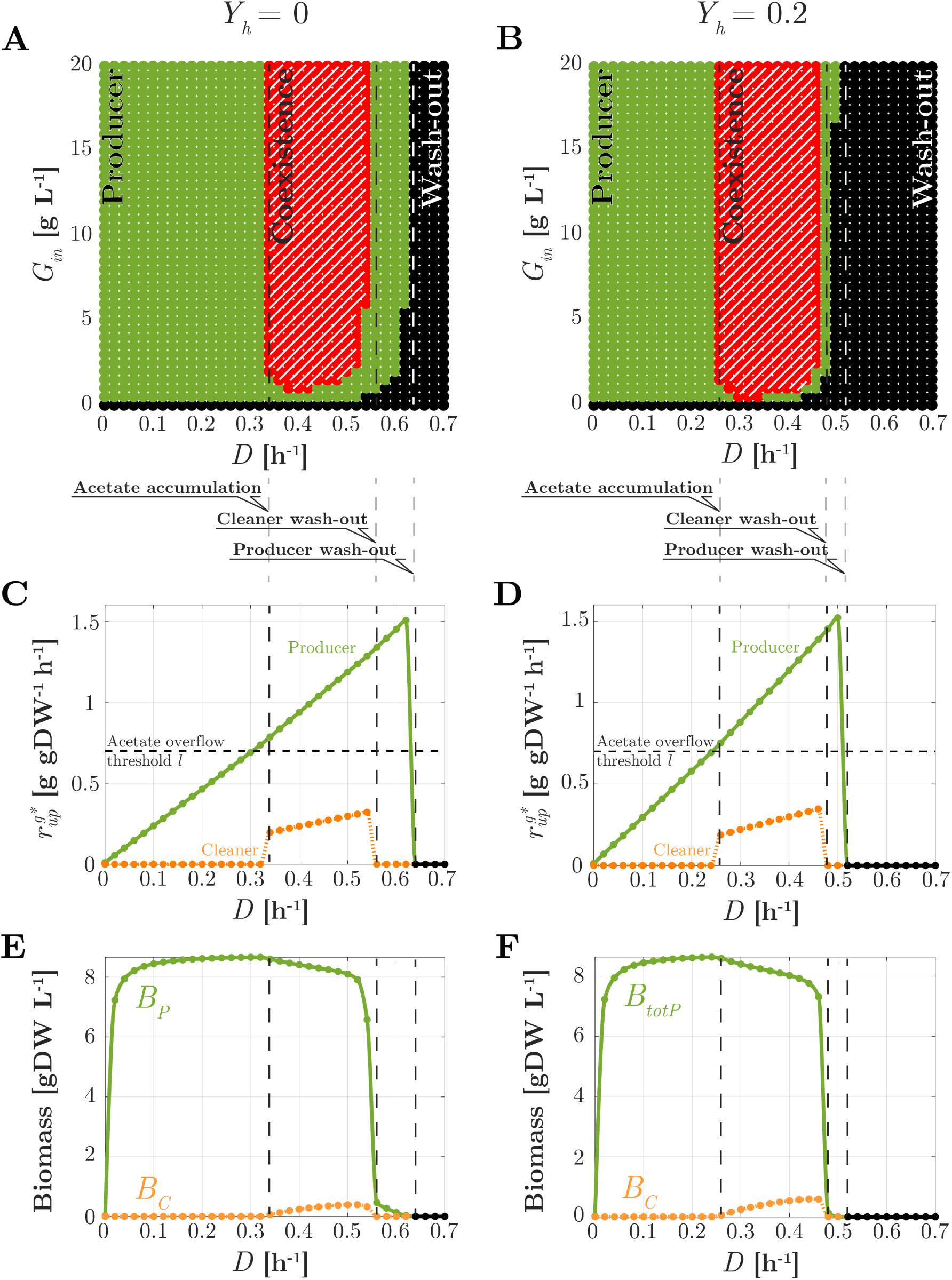
Steady-state analysis of the consortium in chemostat. Left: *Y*_*h*_ = 0; Right: *Y*_*h*_ = 0.2. (A)-(B) Nature of the unique (stable) steady state as a function of *D* and *G*_*in*_ (hatched red: stable coexistence; green: stable existence of the producer only; black: washout of both strains). (C)-(D) For *G*_*in*_ = 20 g L^−1^, for the producer (green dots with solid lines) and the cleaner (orange dots with dotted lines) strains, steady-state value of glucose uptake rate as a function of *D*; (E)-(F) Idem, for biomass and substrate concentrations. Consortium parameters are as in Table 1. Acetate uptake and overflow rates are shown in S6 Fig.

For small values of *D*, 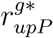 is smaller than the overflow threshold *l*. Notably, in analogy with Eq. 13, and bearing in mind that *Y*_*h*_ = 0, one finds that

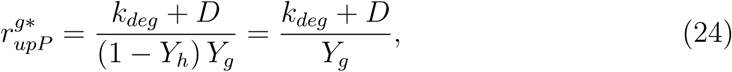

that is, glucose uptake grows linearly with *D* (see Fig. 5C). Absence of acetate overflow impairs cleaner growth due to its inefficient growth on glucose, so that only the producer remains in the bioreactor at this rate.

At a value *D* ≃ 0.34 h^−1^ where Eq. 24 reaches threshold *l*, acetate overflow occurs. In this regime, a comparatively small population of cleaners grows on acetate as soon as the overflow rate is sufficient to sustain growth at the corresponding rate *D*. The resulting cleaner and producer populations balance out in a way that guarantees the precise amount of environmental detoxification to enable growth of the producer at the same rate. The gentle decrease of the producer population size results from utilization of part of the resources to sustain the growth of the cleaner population.

For *D* exceeding 0.55 h^−1^, acetate uptake becomes insufficient to support cleaner growth. Together with the faster utilization of glucose by the producer strain, this results in the washout of the cleaner strain. In absence of acetate scavenging, producer growth at this high rate is only possible at low acetate concentrations, that is, if the population excreting acetate is small. This is witnessed by the sudden drop of 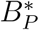.

For *D* above the maximal producer growth rate on glucose, the producer population is also washed out. In view of the dependence of growth rate on glucose concentration via the Monod law (Eq. 10), this threshold depends on *G*_*in*_. Finally, for small values of *G*_*in*_ mildly depending on *D* and typically below 2 g L^−1^, the producer population does not excrete enough acetate to support the cleaner growth at the corresponding rate. Of course, in absence of glucose (*G*_*in*_ = 0), the entire consortium is washed out.

#### Dependence of coexistence on *Y*_*h*_

How is coexistence affected by the synthesis of a heterologous protein? To address this question, we repeat the analysis of the previous section for the case where *Y*_*h*_ = 0.2, which is the value of product yield reported in Table 1. Steady-state results for 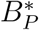 and 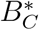 are illustrated in Fig. 5B (a unique stable steady state is again found for every value of *D* and *G*_*in*_, see S4B Fig.). Compared with Fig. 5A, a similar shape of the various coexistence domains is observed. Steady-state rates and concentration profiles qualitatively similar to the case *Y*_*h*_ = 0 are also obtained, as shown in Fig. 5D and 5F. However, quantitative differences in the values of *D* and *G*_*in*_ supporting coexistence are observed.

Coexistence occurs for *D* > 0.25 h^−1^. This value is smaller than the corresponding value for the case where *Y*_*h*_ = 0. This is explained by the fact that, for larger *Y*_*h*_, the necessary acetate overflow to support cleaner existence starts at lower growth (dilution) rates (see the discussion following Eq. 13). For sufficiently large values of *G*_*in*_, coexistence persists up to about *D* = 0.48 h^−1^, whereas the whole consortium is washed out at about *D* ≃ 0.52 h^−1^. Both of these dilution rates are again smaller than their counterparts for *Y*_*h*_ = 0. This is because the metabolic burden associated with *Y*_*h*_ steers away part of the glucose uptake from producer growth, thus decreasing the maximal producer growth rate (Eq. 9) as well as the size of the producer population. The latter implies less acetate excretion and thus a reduction in the maximal cleaner growth rate as well. It is important to note that the interval of dilution rates supporting coexistence, though shifted toward lower values, is comparable to that observed for *Y*_*h*_ = 0. Interestingly, in presence of heterologous protein synthesis, smaller values of *G*_*in*_ (about 1 g L^−1^) are required to observe coexistence at some suitable dilution rate.

We can thus summarize our results with the intriguing observation that, whereas a larger product yield shrinks the survival domain of the producer, it does not shrink the survival domain of the cleaner. In other words, heterologous protein production does not impair coexistence despite causing a metabolic burden on the producer. However, overly large values of *Y*_*H*_ drastically reduce the ability of the producer to grow, to the detriment of the whole community. In fact, it can be seen that when the burden is so heavy that the cleaner outperforms the producer in growth on glucose, regimes where only the cleaner survives are possible (S5 Fig.).

### Coexistence improves productivity but lowers the process yield

We have shown in the previous section that coexistence of the producer and cleaner strains in a chemostat is possible and even favored by a nonzero yield *Y*_*h*_, though at lower dilution (growth) rates. Does coexistence improve performance of the heterologous protein production process? Presence of the cleaner has the potential to favor growth of the producer due to the scavenging of acetate. At the same time, however, growth of the cleaner also consumes glucose, thus taking away resources from the producer. The effects of this resource utilization trade-off on heterologous protein production performance are *a priori* unclear. In the following, we focus on two main questions: When does a consortium achieve the highest production rates? Does the consortium outperform a producer population alone in the synthesis of *H*? As we will see, based on further anaysis of our model in continuous culture conditions in the chemostat (with product yield fixed to *Y*_*h*_ = 0.2), the answer to these questions will unveil further trade-offs between different notions of production performance.

#### The consortium attains highest productivity in a coexistence regime

For the same conditions as in Fig. 5B, Fig. 6A reports a heatmap of the productivity of the consortium as a function of the bioreactor parameters *D* and *G*_*in*_. Productivity is defined as the steady-state rate of outflow of protein *H* through the chemostat effluent (*DH*^*^). Productivity is indeed maximal in a coexistence regime. Highest values are obtained near the maximal dilution rates before the cleaner washout, and for high input glucose concentrations (Fig. 6A). Productivity drops suddenly for dilution rates near the boundary of cleaner washout (*D* ≃ 0.5 h^−1^). This is the result of a sudden decrease of the producer population (Fig. 5F). Productivity is of course zero for larger dilution rates where the producer is also washed out. In summary, the highest productivity is obtained in a regime where coexistence of producer and cleaner is possible. In the next section, we investigate the question whether the consortium is indeed advantageous over a producer strain working in isolation.

**Fig 6.**
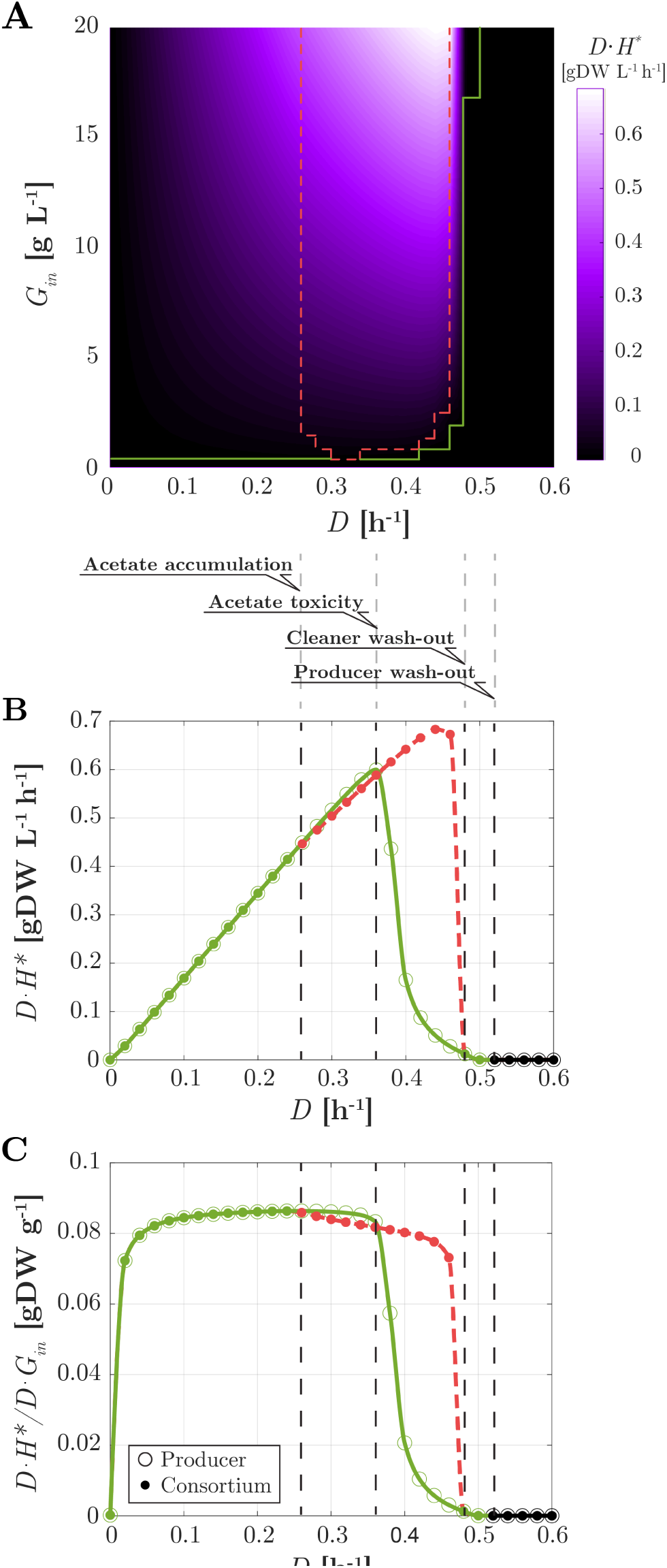
Performance of the heterologous protein production process in chemostat (*Y*_*h*_ = 0.2). (A) Heatmap of the productivity of the consortium (*DH*^*^) as a function of glucose inflow *G*_*in*_ and dilution rate *D*. The boundaries of the domains of coexistence and of existence of the sole producer are reported from Fig. 5B in dashed red and solid green lines, respectively. (B) For *G*_*in*_ = 20 g L^−1^, productivity as a function of *D* for the consortium (filled circles), and for a producer growing in isolation (empty circles). The color code for coexistence (dashed red) or existence of the sole producer (green) is the same as in Fig. 5B. Vertical dashed lines indicate different productivity domains (see discussion in main text). (C) Same as B for the process yield ((*DH*^*^)*/*(*DG*_*in*_)).

#### The consortium attains higher productivity than the producer species alone

Does the consortium outperform a producer population growing alone? Of course, productivity of the former differs from that of the latter only where coexistence is possible, which is also where the consortium attains highest productivity. The question then becomes whether in this region a producer growing in isolation would outperform the consortium.

In Fig. 6B, for the same value of glucose inflow concentration of Fig. 5D and 5F (*G*_*in*_ = 20 g *L*^−1^), we compare productivity of the consortium and of the producer species in isolation as a function of the dilution rate *D*. For the intermediate dilution rates where coexistence is possible, two subdomains can be distinguished. For *D* between 0.26 h^−1^ and 0.36 h^−1^, productivity grows with *D* for both the consortium and the sole producer. However, since part of the resources are devoted to maintain the cleaner species, the producer species alone outperforms the consortium. For values of *D* between 0.36 h^−1^ and 0.48 h^−1^, because of the larger overflow at higher growth rates, the toxic effect of acetate becomes dominant. In absence of the cleaner, this results in a sudden drop of productivity due to a reduced size of the producer population (see S7 Fig.). For the consortium instead, the gentle reduction in producer biomass is overcompensated by the increase of *D*. The net result is an increase in productivity (Fig. 6B), up to the dilution rates where the producer biomass washes out. Crucially, the maximum value of the consortium productivity (about 0.7 g L^−1^ h^−1^ at *D* ≃ 0.44 h^−1^) is about 14% larger than that of the producer alone (about 0.6 g L^−1^ h^−1^ at *D* ≃ 0.36 h^−1^). In summary, the consortium outperforms the producer species alone in terms of productivity. As seen before, maximal productivity is obtained for high values of *G*_*in*_. It is worth noting that the maximum is obtained at a dilution rate strictly smaller than the maximal rate supporting coexistence. For applications, this warrants persistence of the cleaner in spite of fluctuations of the system and modelling inaccuracies. Results along the same lines are obtained for all other values of *G*_*in*_ supporting coexistence.

#### Highest productivity corresponds to a smaller process yield

How efficient is the conversion of substrate into product by the consortium? How does this relate with productivity? To address this point we computed the yield of the process, defined as

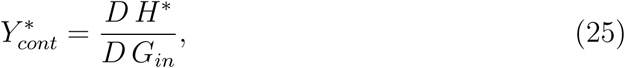

that is, product outflow rate versus substrate inflow rate at steady-state. Fig. 6C shows the yield as a function of *D* for the same value of *G*_*in*_ as in Fig. 6B. Note that, since 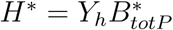 (see *Methods*), it holds that 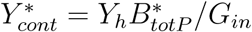. Therefore, the yield of the consortium is proportional to the producer biomass (compare Fig. 6C with Fig. 5F). The same holds for the producer alone. The highest yield is found at the highest dilution rate before overflow (*D* ≃ 0.26 h^−1^), thus in particular, in absence of cleaner. Intuitively, this is because at that point, the largest producer biomass is obtained before resources are partly redirected into acetate overflow. For the consortium, this overflow sustains the cleaner growth. At rates above *D* ≃ 0.26 h^−1^, for the producer alone, biomass loss is initially less pronounced than in the consortium, since no other species consumes glucose. However, at higher dilution rates, this results in strong acetate accumulation, whence a sharp loss of biomass and yield. Compared with Fig. 6B one notices that, for the consortium, maximizing productivity is paid for in terms of a decreased yield. This trade-off takes place because maximal productivity is achieved in presence of the cleaner, that is, in presence of acetate overflow, which subtracts part of the available glucose from the protein production process. Utilisation of glucose is made even less efficient by the glucose uptake of the cleaner, which subtracts further resources from protein production. From a qualitative viewpoint, however, the same trade-off is encountered for a hypothetical cleaner strain without residual glucose uptake (see S8 Fig.).

### Dynamical model and protein production in fed-batch

The productivity of the community is expressed as the rate at which biomass and thus the protein of interest flow out from a bioreactor operating in continuous mode. From a methodological point of view, continuous culture in a chemostat has the advantage of allowing productivity and yield to be analyzed under well-defined steady-state conditions. However, in many biotechnological applications today, fed-batch cultures are still the standard mode of operation [55]. Moreover, the high biomass densities reached during fed-batch operation make acetate accumulation an even more pressing problem. We have therefore extended the analysis of possible productivity gains of the synthetic consortium to fed-batch scenarios.

To simplify the analysis and focus on production performance, we analyze the model under the following assumptions: (i) glucose concentration in the bioreactor, *G*, is kept constant by a suitable time-varying input flow; (ii) there is no outflow; (iii) the total volume in the bioreactor is constant. These assumptions are reasonable over a suitable finite time-horizon and for large ratios between bioreactor volume and input volume. The design of the (time-varying) control input parameters *D* and *G*_*in*_ to meet these assumptions is beyond the scope of this paper. For the consortium, the model describing this scenario is provided by Eqs. 18–21, with *G* fixed and *D* = 0 h^−1^. A similar adaptation holds for the model with producer only.

How does production performance of a consortium and of a producer alone compare in fed-batch? What are the trade-offs involved in this case? We illustrate simulation results for *G* = 20 g L^−1^, also referring to more general theoretical results reported in the *Methods* section. Fig. 7A shows dynamical simulation results for the producer alone and for the consortium with the cleaner. Of course in this case, biomasses do not reach a steady state. As apparent, they rather settle into an exponential growth regime, which they maintain as long as *G* is kept constant. We can thus refer to constant proportions and constant rates in exponential growth (indicated with a superscript “^+^”) to investigate productivity and yield.

**Fig 7.**
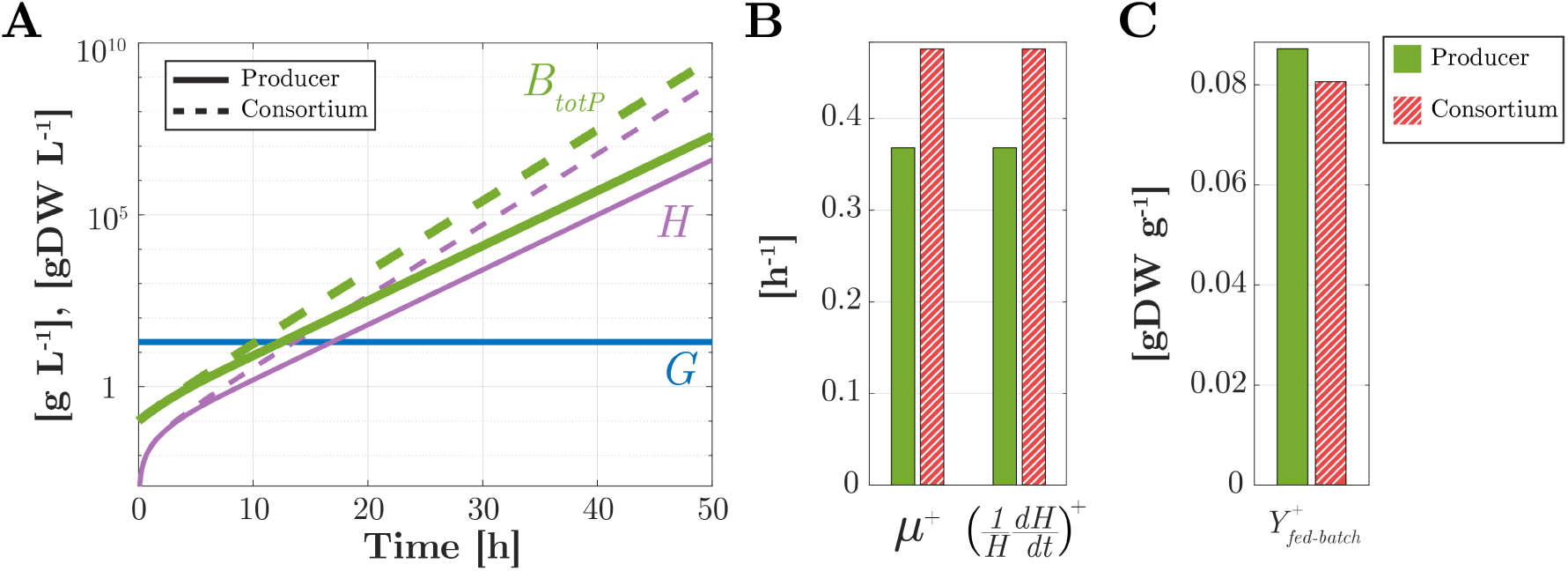
Protein production dynamics in fed-batch. (A) Time evolution in log scale of the concentrations of the total producer biomass (*B*_*tot*_, *B*_*totP*_, thick green) and of the product (*H*, thin purple) for the consortium (dashed lines) and a producer population alone (solid lines) at a fixed environmental concentration of glucose *G* = 20 g L^−1^ (horizontal blue line) (B) Growth rate and rate of increase of *H* in exponential growth, for the consortium (hatched red) and the producer alone (green). (C) Idem for the process yield in exponential growth.

In exponential growth, for both the producer alone and the consortium, the growth rate *µ*^+^ is constant and equal to the constant rate of increase of *H*, that is, (*d* log *H/dt*)^+^ = (*H*^−1^ *dH/dt*)^+^. Moreover, the ratio between the product concentration and the total producer biomass concentration at any point of the exponential growth regime is precisely equal to *Y*_*h*_ (see Fig. 7A and 7B and also *Methods*). However, the rate of increase of *H* is around 0.48 h^−1^ for the consortium and only 0.37 h^−1^ for the producer alone (Fig. 7B). In view of the exponential increase of *H*, it is reasonable to take the rate of increase of *H* as a productivity index. By this criterion, for a fixed environmental glucose concentration, we conclude from the above that the consortium ensures higher productivity of *H*.

How efficient is the conversion of glucose into the protein product? Because of the assumed constant concentration *G*, at any time, the glucose uptake rate by the entire consortium, 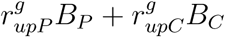, is equal to the rate of glucose supply to the bioreactor. One can thus define the instantaneous yield of the consortium production process in fed-batch as

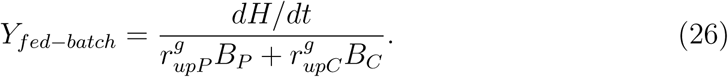

The same index can be defined for the producer alone (where *B*_*C*_ = 0). In exponential growth, this yield reaches a constant value 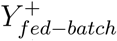, which is equal to 0.087 for the producer alone and 0.08 for the consortium (Fig. 7C). Thus, the yield of the producer alone is found to be greater than that of the consortium.

In summary, similar to the case of a chemostat, we observe a trade-off between productivity and yield of a protein production process in fed-batch. Again, due to the resources consumed in the growth of a subsidiary strain, greater productivity of the consortium is paid for in terms of smaller yield compared to a homogeneous population of producers only.

## Discussion

Synthetic microbial consortia present a promising avenue for a variety of bioengineering applications [1, 3, 5–7]. We are interested in the conditions that allow a synthetic consortium to express a heterologous protein with higher productivity than a single species, as well as the underlying trade-offs shaping these conditions.

As a concrete example, we have chosen a prototypical consortium of a protein producer strain and an acetate cleaner strain of *E. coli* (Fig. 1), a consortium that was experimentally constructed recently [20]. In comparison with the latter consortium, we have envisioned a slightly different design of our cleaner strain. First, while removing the preferential glucose uptake system PTS, we have assumed that secondary glucose assimilation pathways remain functional, which reduces but not completely eliminates growth on glucose of the acetate cleaner strain [52], contrary to the implemented consortium. Second, in order to increase the maximal growth rate on acetate, we have included in the design of the cleaner strain a plasmid for the overexpression of Acs, an enzyme for the irreversible conversion of acetate to acetyl-CoA, as suggested by previous work [53]. The addition of the plasmid facilitates the calibration of the acetate cleaner strain, as it allows reuse of the parameter values for acetate metabolism estimated for the wild-type strain (see *Methods*). Despite these differences, our model reproduces the most important conclusion of Bernstein *et al.* [20], namely that for dilution rates supporting coexistence, the consortium reaches a higher biomass concentration than the wild-type strain alone (S7 Fig.).

A key challenge for constructing synthetic consortia is to guide their design by mathematical modeling and analysis, before the actual implementation *in vivo*. Along these lines, we developed a quantitatively predictive model of the consortium, accounting for a range of growth phenotypes (Fig. 2). Like in other work on the mathematical modeling of community dynamics [28–33], we have developed coarse-grained models of the physiology of the individual species in the consortium, which requires making stark simplying assumptions (see [24, 27] for other perspectives, based on genome-wide flux balance models). Despite the simplifications, however, a critical requirement to assess the potential for productivity gains is the ability of the model to qualitatively and quantitatively reproduce nutrient production and consumption patterns, as well as to predict the growth rates of individual species. In the case of the *E. coli* consortium studied here, this requires to include a variety of regulatory phenomena (threshold for acetate overflow, acetate toxicity, carbon catabolite repression, growth-independent maintenance, …), like in [45, 46], that have not or only partially been accounted for in previous models of this and structurally similar consortia. The resulting model, with only a single biomass variable for each species, is able to quantitatively reproduce complex growth phenotypes like diauxic growth (Fig. 3A) the effect of energy dissipation on acetate overflow (Fig. 3B), and the effect of glucose uptake mutants on growth and acetate overflow (Fig. 3C-D). The analysis of the model of the synthetic consortium shows that, over the entire range of dilution rates, the system has a unique stable steady state (Fig. 5). These numerical results were confirmed by explicit computation and analysis of the roots of the multinomial equations (that is, polynomial equations in several variables) characterizing the steady states of the consortium (S4 Fig.). Another model of the same consortium, very similar in the level of detail of the description of the community dynamics [31], predicted the possibility of multiple stable steady states, one corresponding to coexistence of the producer and the cleaner strains, and the other to existence of the producer strain alone. The property of multistability was found to rely on the non-monotonicity of the function describing the dependence of the growth rate on the acetate concentration in the medium, when acetate is used as the sole carbon source [31]. The maximal growth rate is thus reached for intermediate acetate concentrations instead of being monotonically approached for high acetate concentrations, like in Eq. 11. The Monod-like function for acetate uptake used in this study was motivated by experimental data describing a monotonic increase of the growth rate with increasing acetate concentration, over the relevant range considered here [16, 56]. The non-monotonicity of the function used by Harvey *et al.*, however, is also supported by experimental data (Tomas Gedeon, personal communication). The discrepancy between the studies might be due to the use of different *E. coli* strains, different media compositions, or other details of the experimental procedures.

The model predicts that coexistence of the protein producer and acetate cleaner strains is possible over a range of dilution rates (Fig. 5). While coexistence in natural and synthetic ecosystems has been studied in previous work [30, 31, 33], and is an obvious prerequisite for the possibility of increasing productivity by a consortium, the observation that this dependency works in both directions, in that the production of a heterologous protein affects coexistence as well, is a novel finding of this study. In particular, the analysis of the model shows that the coexistence region is shifted towards lower dilution rates (Fig. 5), as a consequence of the metabolic load associated with heterologous protein production. The effect of metabolic load on microbial growth has been well-studied and is an active subject of research in synthetic biology [37, 57], where feedback strategies for avoiding the deleterious consequences of metabolic overloading have been developed [58]. Our results show that questions on the relation between growth and metabolic load appear on the community level as well, and are important for assessing the productivity of synthetic consortia. Whereas the biomass concentration attained, in the absence of the production of a heterologous protein, has sometimes been used as a proxy for the productivity of a consortium [59], our analysis shows that subtle effects involving feedback from metabolic load to growth may be at work. For example, in our system maximal productivity is obtained for a dilution rate where the total biomass concentration is suboptimal.

One of the main conclusions from this study is that coexistence of the producer and the cleaner strains may lead to a productivity gain (Fig. 6). Coexistence alone is not enough to ensure this gain though, as the latter ultimately depends on a trade-off. For lower dilution rates in the coexistence region, the toxic effect of acetate is very limited and thus investing part of the substrate in the growth of non-productive acetate scavengers is not profitable, since it has the sole effect of reducing the producer population. Indeed, despite growth inhibition by acetate, the producer strain in monoculture manages to reach a concentration that is sufficient for supporting a higher protein production rate than in coculture. For higher dilution rates in the coexistence region, however, the toxic effect of acetate becomes so strong that the producer concentration, and thus productivity, plummet when the producer is grown without the cleaner. The investment of resources necessary to sustain the cleaner population pays off in this case: removing acetate from the medium allows heterologous protein production at a rate that more than compensates the utilization of glucose by the cleaner. Note that this trade-off involves a subtle balance between growth of the producer and cleaner, acetate overflow, acetate inhibition, and protein production, that is, nonlinear dynamical processes with feedback operating at the level of the individual strains and their interactions. Therefore, predicting the productivity associated with specific bioreactor control parameters would have been extremely difficult, if not impossible, to achieve without the use of quantitative dynamical models of the type developed here.

Higher productivity in the consortia comes at the price of a lower process yield, that is, the outflow rate of heterologous protein divided by the inflow rate of substrate decreases as the productivity increases (Fig. 6). This is basically another manifestation of the trade-off discussed above. Higher productivity requires a higher dilution rate, and thus more acetate overflow. In particular, maximal productivity is attained in presence of the cleaner thanks to its consumption of acetate, which allows the producer to grow at rates where acetate concentration in the medium would otherwise have reached toxic levels. However, the acetate overflow that sustains the cleaner growth at such high rates subtracts significant amounts of glucose from conversion into the target protein, which results in yield loss. The rate-yield trade-off is well-known in metabolic engineering [39] and has been extensively studied in microbial physiology [40], mostly in the context of single species or strains. We here show that this trade-off also appears on the community level, when weighing the costs of sustaining a cleaner population against the benefits of increased protein production. The rate-yield trade-off, as well as the higher productivity of the consortium of protein producers and acetate cleaners, is not only predicted in continuous culture in a chemostat, but also in fed-batch conditions (Fig. 7).

While our modeling study has focused on a specific synthetic consortium of protein producers and acetate cleaners (Fig. 1), the conclusions of our study may be more general and carry over to structurally similar synthetic consortia reported in the literature [60]. An example is the consortium consisting of *Ralstonia eutropha* and a Δ*manA* mutant of *Streptomyces coelicor* for the production of biodiesel [61]. *S. coelicor* Δ*manA* produces fatty acid methyl esters, while secreting acetate and citrate into the medium, resulting in growth inhibition and a lower pH. *R. eutropha* utilizes the fermentation acids, thus promoting growth of the producer strain and increasing pH. Another example is the consortium consisting of a *S. cerevisiae* strain carrying an enzyme for the production of methyl halides and the cellulolytic bacterium *Actinotalea fermentans* [62]. The latter ferments cellulose to acetate and ethanol, which the former utilizes for growth and methyl halide production. The consumption of the fermentation by-products by the yeast strain lifts their inhibitory effect on *A. fermentans* growth. Note that in this consortium, the cleaner produces the metabolite of interest, whereas the producer provides the growth substrates. We expect that our conclusions, for example, that metabolic load influences producer and cleaner coexistence, or that higher productivity comes at the price of lower yield, will also apply, with appropriate qualifications, in these and other cases.

In the current analysis of the consortium we have assumed that the only control inputs we dispose of are the dilution rate and the glucose concentration in the inflow, and we have determined productivity and yield only at steady state. The question under which conditions the consortium can achieve higher productivity can also be addressed with a larger range of control inputs, such as the possibility to change *Y*_*H*_ and *k*_*Acs*_, to give but two examples. Moreover, appropriate control laws may be needed to stabilize the consortium against fluctuations of the bioreactor operating conditions, modelling errors, *etc.*, or, in case of multistability, to drive the system to the appropriate steady state. When considering the consortium from a dynamical perspective, one can formulate optimal control problems for the maximization of heterologous protein production in finite time and/or dynamical conditions, instead of maximizing productivity at steady state [63]. The actual implementation of the optimal dynamical control schemes will require feedback control of the community composition and the functioning of the individual species, in the line of recent proposals in the literature [64, 65]. This suggests exciting problems at the interface of control theory, synthetic biology, and biotechnology.

## Methods

### Parameter estimation and identifiability analysis

The model of the producer strain is given by Eqs 1-12 and has 14 parameters, most of which were estimated from a recently published data set [13]. The data concern steady-state exponential growth in batch of the *E. coli* wild-type strain K-12 MG1655 in minimal medium with (i) glucose as the sole carbon source, (ii) acetate as the sole carbon source, and (iii) glucose with different concentrations of acetate added. The growth conditions of the experimental data set (wild-type strain in batch) imply that *Y*_*h*_ and *D* must be set equal to 0.

The data set was completed with measurements of the degradation constant *k*_*deg*_ [66, 67], *the threshold for acetate overflow l* [68] *and the half-maximal saturation constants K*_*g*_ [69] *and K*_*a*_ [70]. *The value of k*_*deg*_ was computed from experimental quantification of the maintenance coefficient *C*_*m*_ [71] *in conditions similar to ours, using the relation k*_*deg*_ = *C*_*m*_ *Y*_*g*_. The value of *K*_*a*_ was chosen so as to match the measured constant for the Pta-AckA pathway, the main uptake pathway for the range of acetate concentrations considered here [12].

The estimation of the other parameters from the experimental data set was carried out in two steps. First, the parameters *k*_*g*_, *k*_*over*_, *k*_*a*_, *Y*_*g*_, and *Y*_*a*_ were fixed by analytically solving the reduced systems of equations at steady state and using the measured rates (i) in the absence of acetate uptake, and (ii) in the absence of glucose and therefore acetate overflow (Fig. 2B). Second, the remaining parameters Θ_*a*_, *n*, Θ_*g*_, and *m* were estimated from the measured rates in case (iii) above, by means of an optimization procedure that minimizes the sum of squared errors between the predictions and measurements of the growth rate (*µ*^+^), the glucose uptake rate 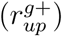, and the net acetate uptake rate 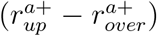 during exponential growth in batch (Fig. 2C). The optimization procedure made use of the `fminsearchbnd` function of Matlab available at mathworks.com/matlabcentral/fileexchange. We evaluated the results by performing an *a-posteriori* identifiability analysis using a procedure analogous to bootstrapping as described before [72], resulting in confidence intervals for the parameter estimates (S1 Fig.).

The parameter values of the calibrated model are summarized in Table 1. There is a good correspondence of the estimated values with other reported values in the literature. For example, the glucose and acetate yield coefficients are in agreement with the measured values of *Y*_*g*_ ≃ 0.44 g gDW^−1^ [73–75] and *Y*_*a*_ ≃ 0.3 g gDW^−1^ [13,46]. The experiments used for model calibration were carried out at pH 7 [13]. High-density fermentation in industrial bioreactors may operate at a lower pH though [76], exacerbating the growth-inhibitory effect of acetate. We therefore used a second data set on the effect of acetate on the growth rate of *E. coli*, carried out at pH 6.4 [16], to adjust the parameter Θ_*a*_ characterizing acetate inhibition. This decreased the value of Θ_*a*_ from 3.5 g L^−1^ to 0.52 g L^−1^ (S2 Fig.).

Unless stated otherwise, the value of the product yield *Y*_*h*_ has been set to 0.2 in the simulations, meaning that 20% of the protein synthesis capacity has been deviated to the production of heterologous protein, a level that can routinely be reached [77].

The model of the cleaner strain is a modified version of that of the producer strain. The cleaner does not produce the heterologous protein and two genetic modifications change the rate equations (Eq. 14 and Eq. 16). This leads to three additional parameters (Table 1). We set *k*_Δ*PTS*_ = *k*_*g*_*/*4 [52] and *K*_*Acs*_ = 0.012 g L^−1^ [13]. Moreover, we chose mild overexpression for the Acs enzyme, *k*_*Acs*_ = 1.5 *k*_*a*_, reflecting the observation that overexpression of Acs in minimal medium with acetate as the sole carbon source leads to faster growth [53] (in the K-12 MG1655 strain, contrary to the W strain [78]).

### Derivation of yield equations

The *Results* section features a number of expressions showing the relation between the product yield *Y*_*h*_ and other quantities at steady state in continuous culture (Eq. 9) or for a constant growth rate in fed-batch culture. Below we derive these expressions from the model equations.

First, at steady state, when equating Eq. 3 to 0 and dividing by *B*, we find

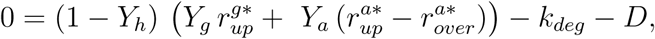

and similarly, when equating Eq. 4 to 0 and dividing by *H*,

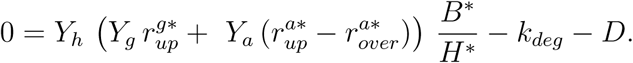

Now, when equating the right-hand sides of these two expressions, and eliminating shared terms on both sides, we are left with

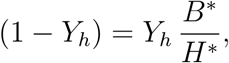

so that with *B*_*tot*_ = *B* + *H*, and solving for *Y*_*h*_, we obtain

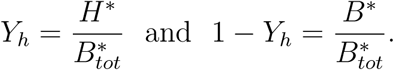

The latter expressions leads to Eq. 9. Note that, due to the equivalence of Eq. 3–4 and of Eq. 19–20, the same relationships hold for the biomass of the producer in the consortium.

Second, we consider the fed-batch case where the bioreactor glucose concentration *g* is set to a constant (assuming suitable glucose inflow) and the rate of the outflow is set to *D* = 0 h^−1^. The relationships we derive again apply to the producer alone or in consortium with the cleaner. The relevant condition to study in this case is exponential growth (see, *e.g.*, Fig. 7A), that is, zero second derivatives of log *B*_*P*_ (*t*) and log *H*(*t*). From Eq. 19, the second derivative of log *B*_*P*_ (*t*) is given by

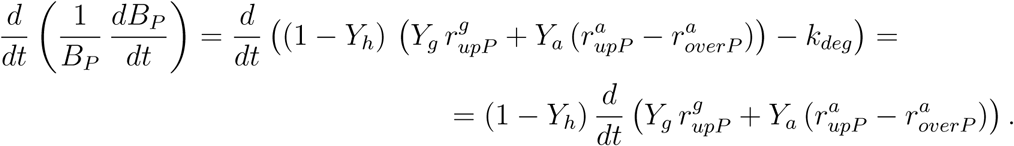

Setting this to zero implies that the sum of rates 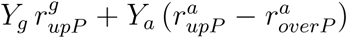 is a constant which we denote by *r*^+^. From Eq. 20, the second derivative of log *H*(*t*) is then given by

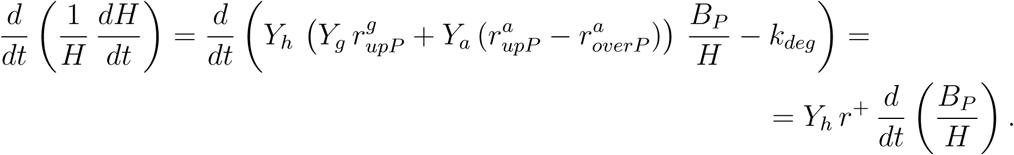

Setting this to zero implies that the ratio *B*_*P*_ */H* is a constant which we denote by (*B*_*P*_ */H*)^+^ (different from above, note that this does not imply that either *B*_*P*_ or *H* is constant). Then log(*B*_*P*_ */H*) is also a constant. By the same relationships employed above, one has

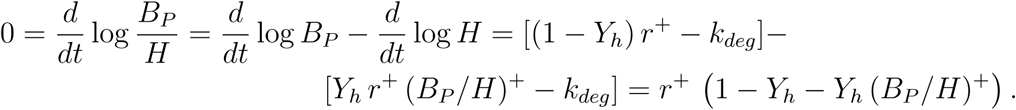

Restricting attention to the nontrivial case where *r*^+^ ≠ 0 (that is *G* > 0), the above implies *Y*_*h*_ = (1 + (*B*_*P*_ */H*)^+^)^−1^. Since (1 + *B*_*P*_ */H*)^−1^ = *H/*(*H* + *B*_*P*_) and 1 − (1 + *B*_*P*_ */H*)^−1^ = *B*_*P*_ */*(*H* + *B*_*P*_) for any *H* and *B*_*P*_, when *B*_*P*_ */H* is constant, *H/B*_*totP*_ and *B/B*_*totP*_ are also constant, given by (*H/B*_*totP*_)^+^ = (1 + (*B*_*P*_ */H*)^+^)^−1^ and (*B/B*_*totP*_)^+^ = 1 − (1 + (*B*_*P*_ */H*)^+^)^−1^. In summary, for this case,

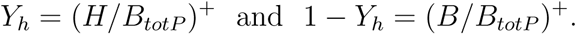

### Computation of steady states

Eqs 17–21, with rate functions defined by Eqs 10–11 and Eqs 14–16, make up an Ordinary Differential Equation (ODE) system with state vector *x* = (*G, A, B*_*P*_, *H, B*_*C*_) taking values in 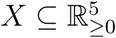. Let us express the system in compact notation as

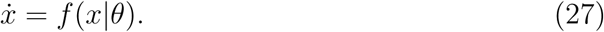

Here *θ* is the vector of parameters that are varied in the study of the system, that is, *θ* = (*D, G*_*in*_, *Y*_*h*_), while the other system parameters are fixed as in Table 1. In order to compute the stable steady states of this system for different values of *θ*, we employ the following two methods.

#### Numerical approach

In this approach, stable steady states of the system are sought by numerical integration of Eq. 27. Recall that, for an assigned initial condition *x*(0) = *x*_0_, a (strictly) stable equilibrium of the dynamical system is given by the asymptotic value of the corresponding solution 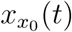, that is, by 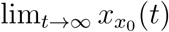, as long as this exists, it is unique and finite. In practice we obtain this by integrating Eq. 27 from *x*_0_ = 0 by the Matlab ODE solver *ode15s* over a sufficiently long time horizon (*T* = 10^7^ h). This procedure is computationally efficient and allows for fast exploration of the system over large sets of values of *θ*. However, for any given *θ*, if the system has several stable steady states, then the steady state eventually reached will depend on *x*_0_. This has motivated the parallel use of a second, algebraic approach.

#### Algebraic approach

For any given *θ*, the possible stable steady states of Eq. 27 may as well be sought as the real, nonnegative solutions in *x* of the equation

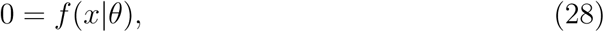

such that the Jacobian matrix *F* (*x*|*θ*) = *∂f* (*x*|*θ*)*/∂x* is strictly stable (that is, all its eigenvalues have negative real part). In the interest of clarity and simplicity, in what follows we drop *θ* from the notation, and instead write rates as functions of *x*. It can be verified by inspection of Eqs 10–11 and Eqs 14–16 that *f* (*x*) is a piecewise rational function. Indeed, the piecewise linear structure of Eq. 12 and Eq. 15 induces a finite partitioning of the state space *X* into the four subsets

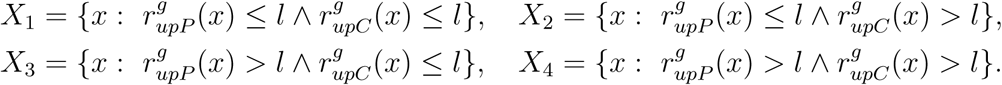

Over each *X*_*i*_, *f* (*x*) = *N*_*i*_(*x*)*/D*_*i*_(*x*), where *N*_*i*_(*x*) and *D*_*i*_(*x*) are two multinomial functions determined by the relevant form of Eq. 12 and Eq. 15 (for instance, the rate conditions that define *X*_2_ imply that Eq. 12 is equal to 0 and Eq. 15 is equal to 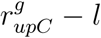 in that domain). For every *i*, the solutions of Eq. 28 belonging to *X*_*i*_ correspond to the roots in *X*_*i*_ of *N*_*i*_(*x*). Out of these solutions, for *i* = 1, …, 4, the steady states sought are the real nonnegative solutions such that *F* (*x*) is strictly stable.

Multinomial root finding is a well explored problem and specialized tools for enumerating all solutions exist [79]. In addition, both *F* (*x*) and the *N*_*i*_(*x*) can be computed explicitly or symbolically at little cost. This suggests the following algorithm:

1. Compute *N*_*i*_(*x*), with *i* = 1, …, 4, and *F* (*x*); For *i* = 1, …, 4:
2. Find the real nonnegative solutions of *N*_*i*_(*x*) = 0;
3. Record solution *x* as a system stable steady state if and only if *x* ∈ *X*_*i*_ and *F* (*x*) is strictly stable.

In our implementation, which is optimized by suitable simplifications for the exploration of steady states with *B*_*P*_ = 0 or *B*_*C*_ = 0, symbolic calculations (step 1) are performed in Wolfram Mathematica, while multinomial root finding (step 2) is performed by the Matlab (numerical) routine *psolve* available at NAClab [80, 81]. In practice, step 3 simply amounts to checking that the evaluation of 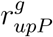 and 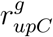 at the candidate solution *x* agrees with the rate conditions defining *X*_*i*_, thus avoiding the explicit calculation of the latter. In view of the complexity of the multinomials *N*_*i*_(*x*), which comprise terms of order up to 33, this method is computationally intensive, yet it returns all possible steady states within the precision of the routine *psolve*.

## Acknowledgments

This work was supported in part by the Inria IPL project CoSy and by the French national funding agency under project Maximic (ANR-17-CE40-0024). The authors thank Johannes Geiselmann (Université Grenoble-Alpes) and Carlos Martinez Von Dossow (Inria Sophia Antipolis – Méditerranée) for profitable discussions and careful reading of the manuscript.

## Supporting information

### S1 Fig Results of the *a-posteriori* parameter identifiability analysis^*^

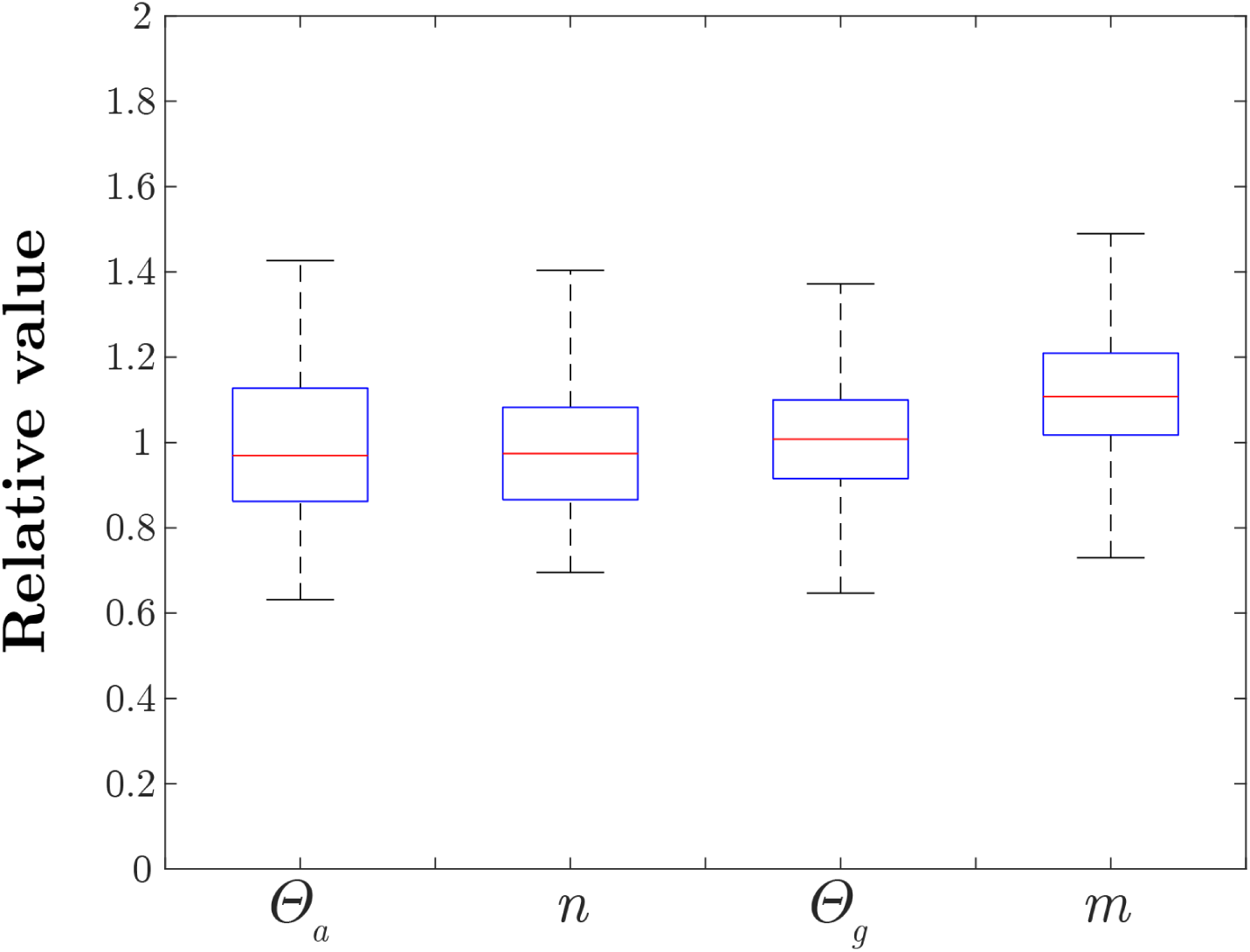

Result of the *a-posteriori* identifiability analysis using a procedure analogous to bootstrapping as described in [1]. We generated 1000 new datasets from the experimental dataset of Fig. 2C [2] by adding to each one of the estimated data points a quantity randomly sampled from the set of residuals obtained by comparing the model predictions and the experimental data. The resulting bootstrap datasets were used to fit the model and obtain new values for Θ_*a*_, *n*, Θ_*g*_ and *m*. The box plot shows the statistic resulting from this procedure. Data are normalized with respect to the estimated value reported in Table 1. The central line of each box represents the median, the bottom and the top edges correspond to the lower and upper quartiles, respectively, and the whiskers extend from 1.5 IQR (interquartile range) below the lower quartile to 1.5 IQR above the upper quartile (see the *boxplot* function in Matlab).

## S2 Fig Model predictions for the producer strain^*^

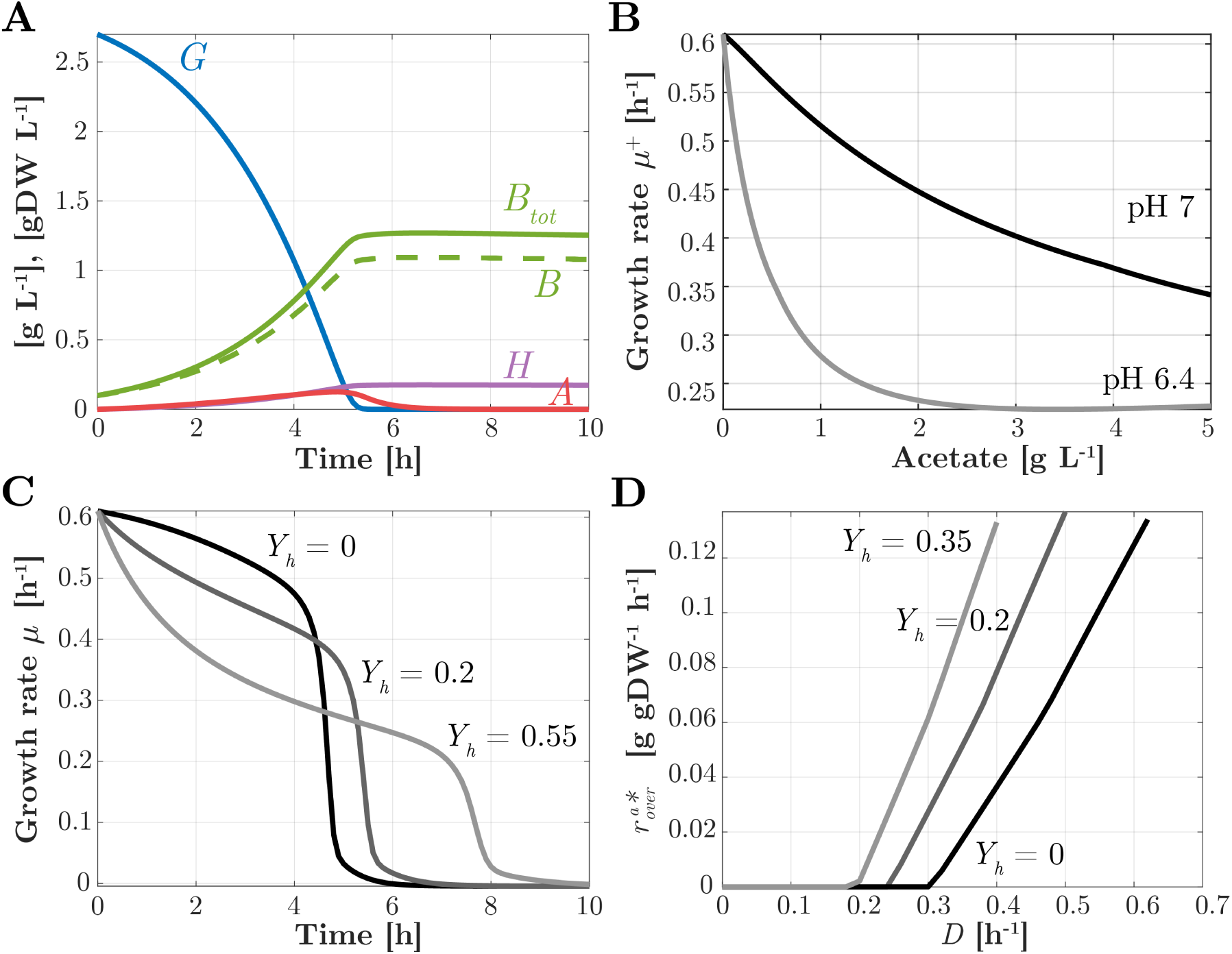

Model predictions for the producer strain. (A) Time evolution of the concentration of glucose *G* (blue curve), acetate *A* (red curve), heterologous protein concentration *H* (violet curve), autocatalytic biomass *B* (green dashed curve) and total biomass *B*_*tot*_ = *B* +*H* (green solid curve) in a bioreactor operating in batch mode, with initial concentrations of 2.7 g L^−1^ of glucose and 0.1 g L^−1^ of biomass. (B) Effect of increasing concentrations of acetate in the medium on the exponential growth rate of the bacteria, in the same conditions as in panel A, with the acetate inhibition coefficient Θ_*a*_ set equal to 3.5 g L^−1^ and 0.5 g L^−1^, corresponding to pH 7 and 6.4, respectively. (C) Effect of metabolic load on the time-varying growth rate in the same conditions as in panel A, obtained by overexpressing the heterologous protein *H* for three different yield coefficients *Y*_*h*_ = 0, *Y*_*h*_ = 0.15, and *Y*_*h*_ = 0.55. (D) Effect of overexpression of heterologous protein *H* on the onset of acetate overflow, indicated by the overflow rate 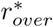 reached at steady state in a bioreactor operating in continuous mode, with *G*_*in*_ = 20 g L^−1^, for three different yield coefficients (*Y*_*h*_ = 0, *Y*_*h*_ = 0.15, *Y*_*h*_ = 0.25), and *G*_*in*_ = 20 g L^−1^.

## S3 Fig Example simulation of the consortium in conditions leading to coexistence^*^

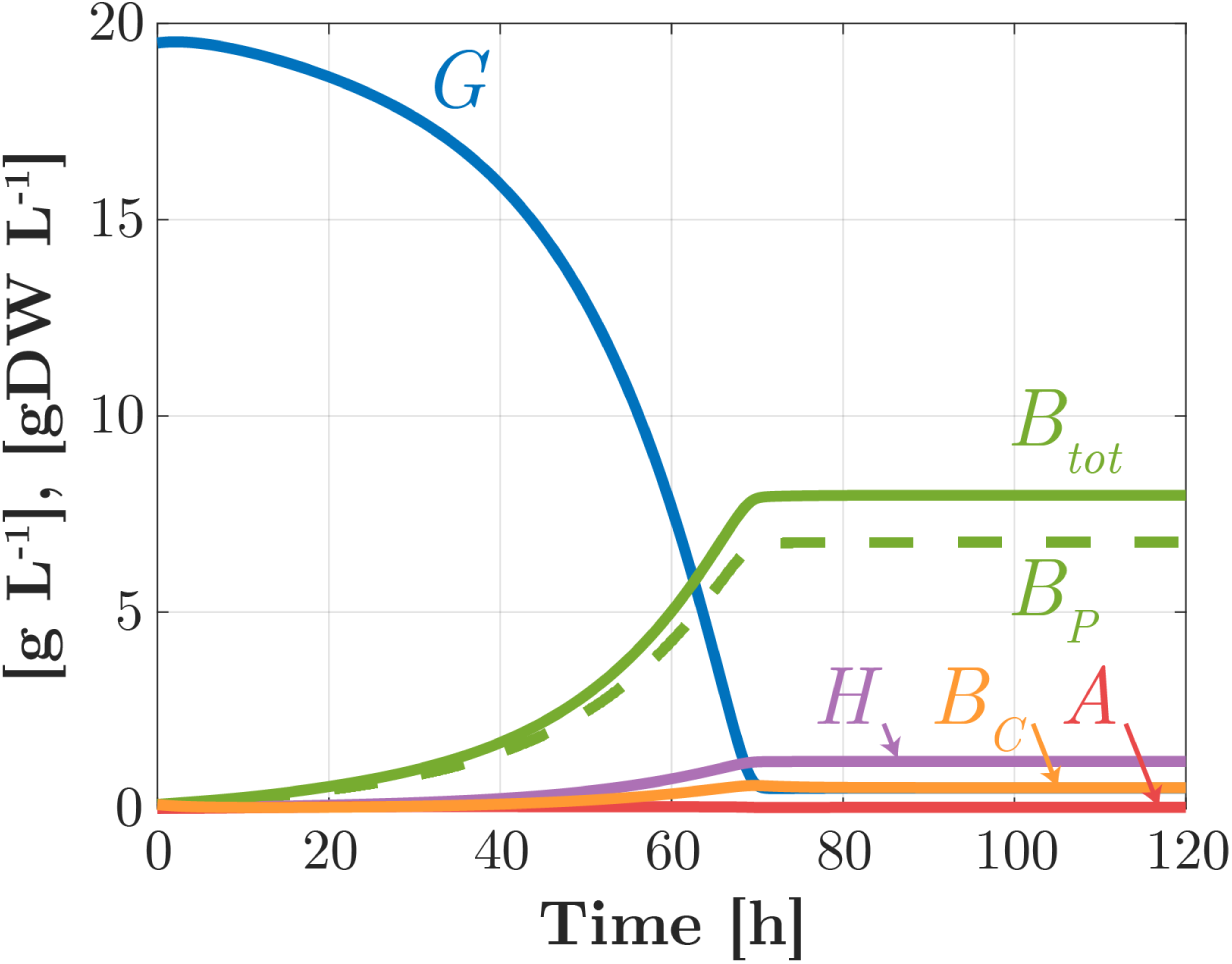

Example simulation of the consortium in conditions leading to coexistence with initial conditions *B*_*P*0_ = 0.1 gDW L^−1^, *B*_*C*0_ = 0.1 gDW L^−1^, *G*0 = 19.5 g L^−1^, and all other initial values set to 0.

## S4 Fig Explicit computation and analysis of the roots of the multinomial equations characterizing the steady states of the consortium^*^

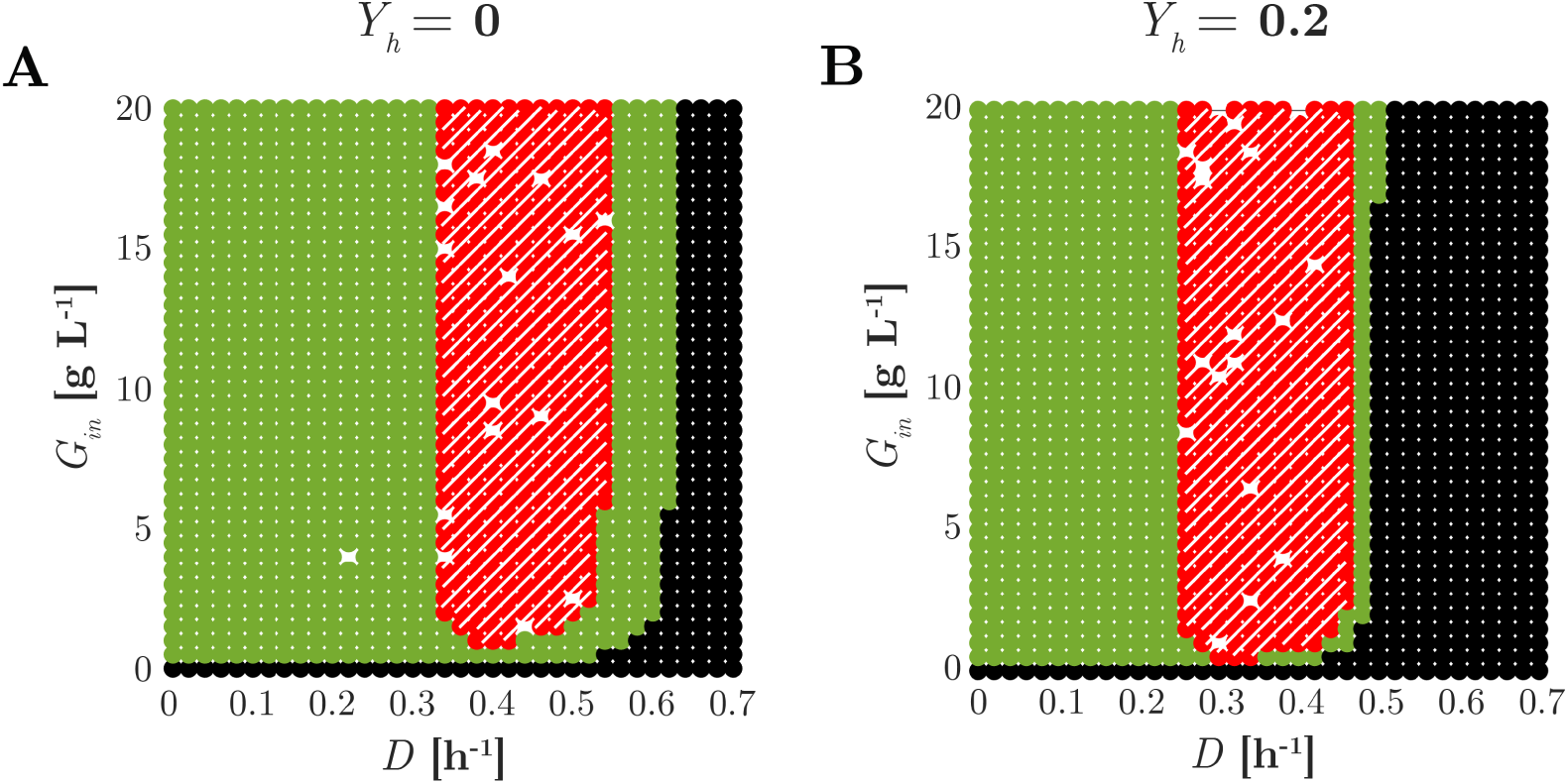

Explicit computation and analysis of the roots of the multinomial equations characterizing the steady states of the consortium obtained as described in *Methods* for *Y*_*h*_ = 0 (A) and *Y*_*h*_ = 0.2 (B). In green, hatched red and black, the region where only the producer, both producer and cleaner, and none of the two strains are present at steady state, respectively. The *psolve* routine of Matlab [1] was not always able to produce the correct solution due to numerical issues, as shown by the sparse blanks.

## S5 Fig Analysis of the unique stable steady state of the consortium, and corresponding productivity heatmap, as a function of *D* and *Y*_*h*_^*^

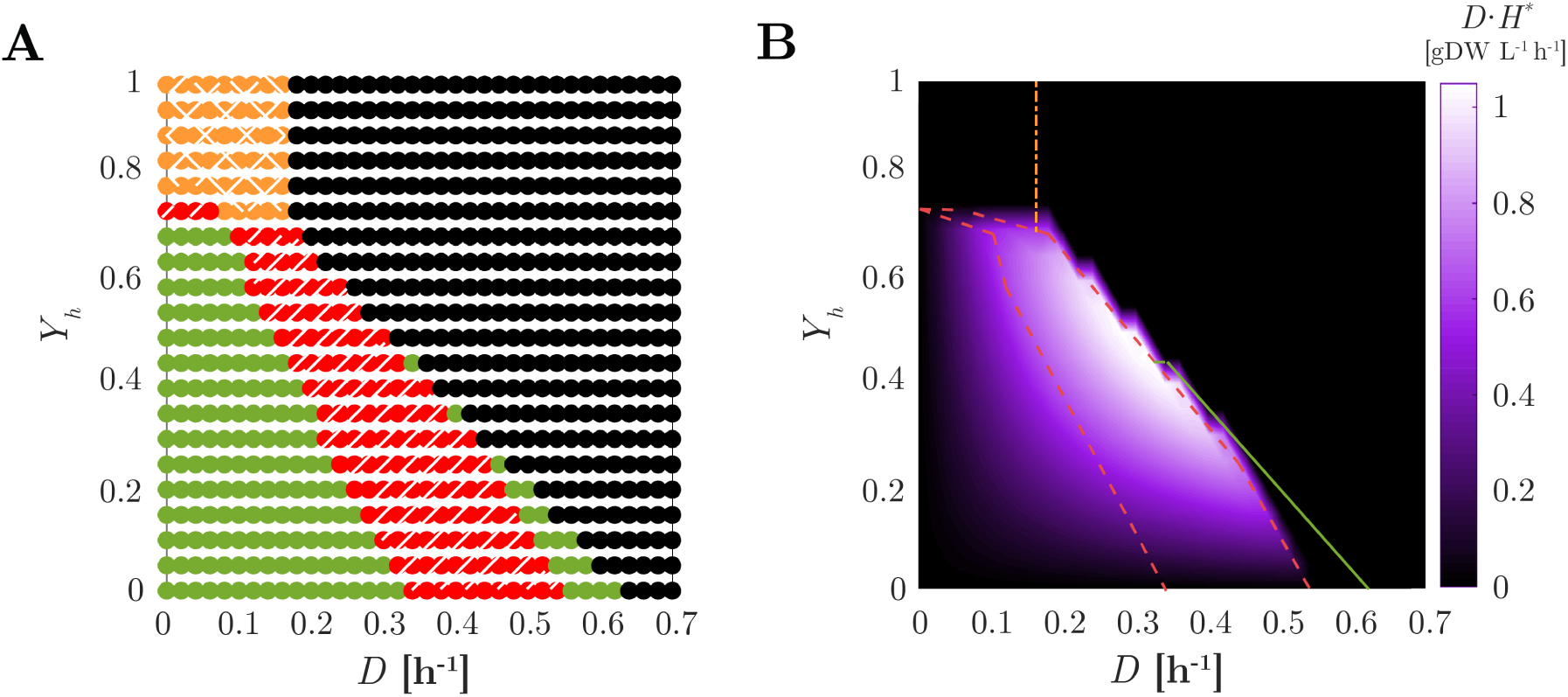

(A) Analysis of the unique stable steady state of the consortium as a function of *D* and *Y*_*h*_. In hatched red: stable coexistence; in green: stable existence of the producer only; in doubly-hatched orange: stable existence of the cleaner only; in black: washout of both strains. (B) Heatmap of the productivity of the consortium (*D H*^*^) as a function of *D* and *Y*_*h*_. The boundaries of the domains of coexistence. of existence of the sole producer, and of the sole cleaner are reported from Fig. S5A in dashed red, solid green, and dahs-dotted orange lines, respectively.

## S6 Fig Acetate overflow rate 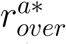 and uptake rate 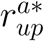 as a function of the dilution rate *D*, for the consortium in steady state in the same conditions as in Fig. 5^*^

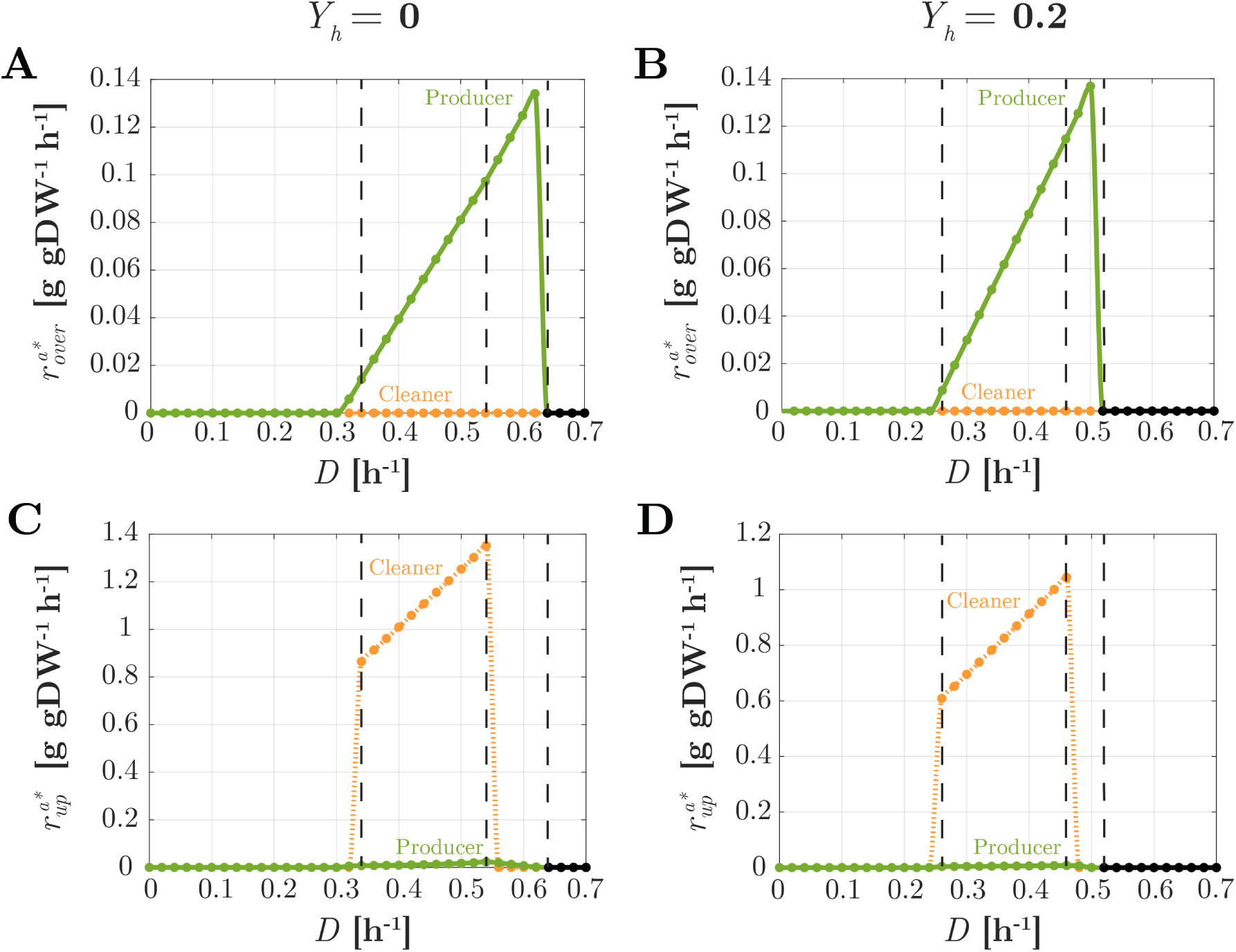

Acetate overflow rate 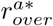 (A)-(B) and acetate uptake rate 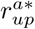 (C)-(D) as a function of the dilution rate *D*, for the consortium in steady state in the same conditions as for Fig. 5 in main text (*G*_*in*_ = 20 g L^−1^). Left panels: *Y*_*h*_ = 0; right panels: *Y*_*h*_ = 0.2. Green dots with solid lines: producer rates; orange dots with dotted lines: cleaner rates. Black denotes absence (washout) of both strains.

## S7 Fig Total steady-state biomass production in chemostat (*Y*_*h*_ = 0.2) as a function of *D* for the consortium and for a producer growing in isolation^*^

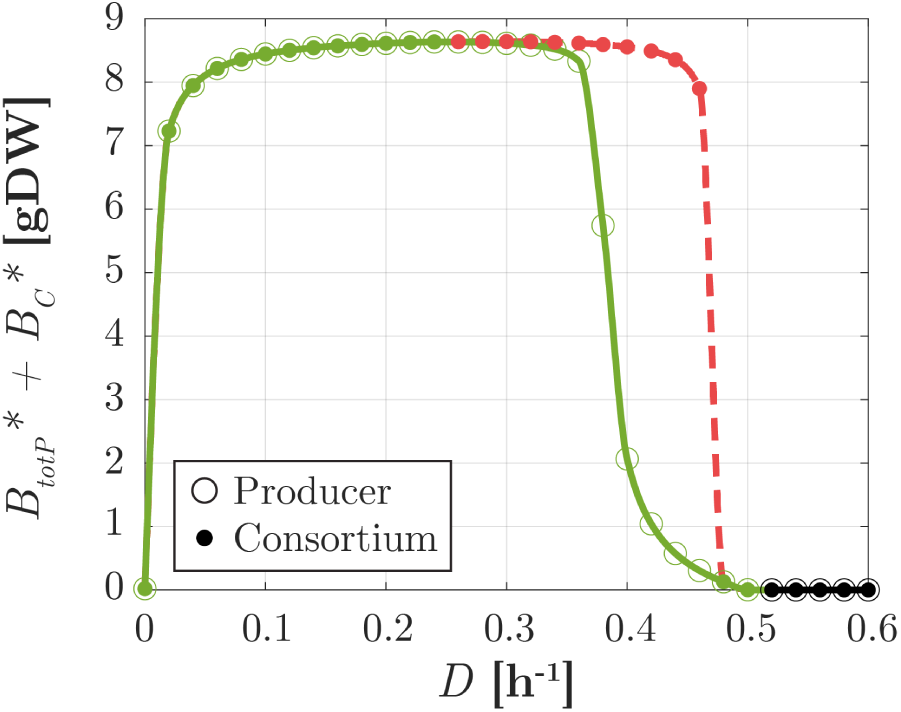

Total steady-state biomass production 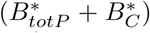 in chemostat (*Y*_*h*_ = 0.20) for *G*_*in*_ = 20 g L^−1^ as a function of *D* for the consortium (filled circles), and for a producer growing in isolation (empty circles), as in Fig. 6. In dashed red: stable coexistence; in solid green: stable existence of the producer only; in black: washout of both strains.

## S8 Fig Performance of the heterologous protein production process in chemostat (*Y*_*h*_ = 0.2), in the case where the cleaner uptake rate of glucose is fixed to 0^*^

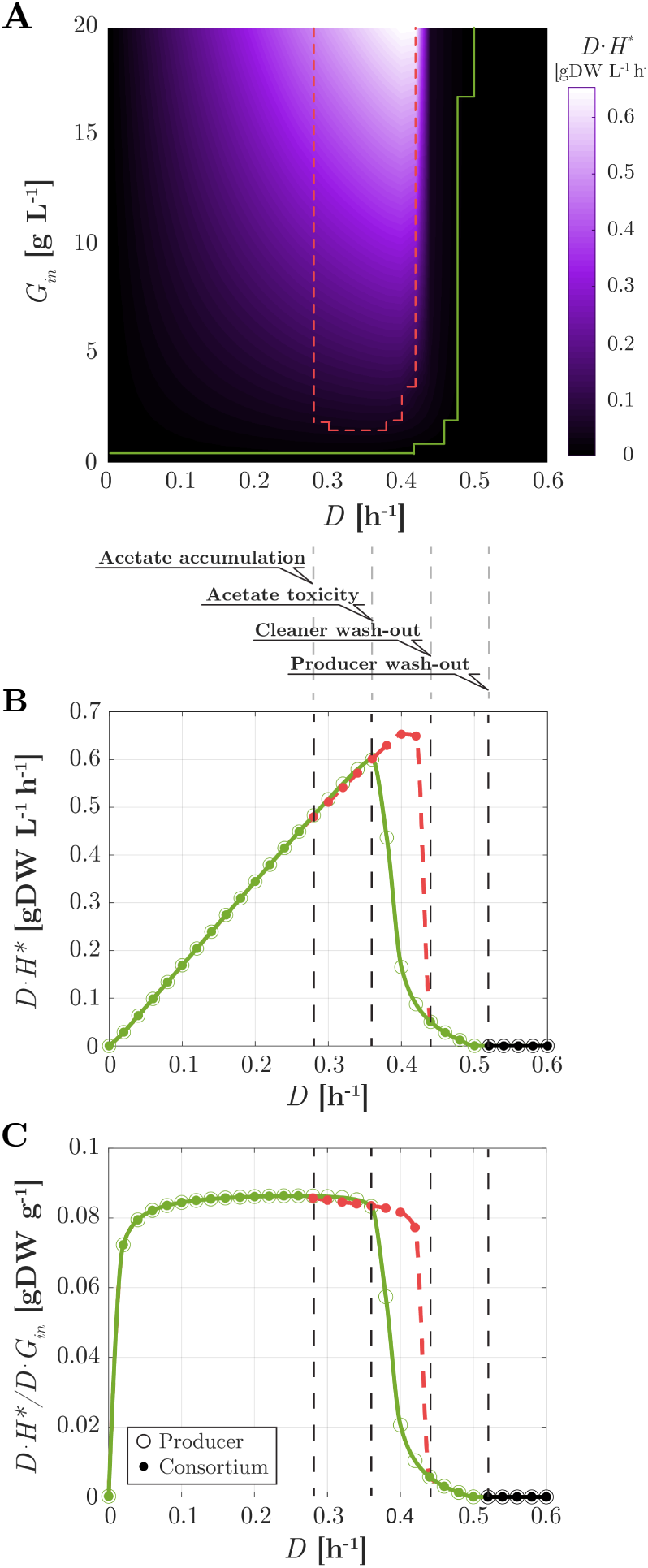

Performance of the heterologous protein production process in chemostat (*Y*_*h*_ = 0.2), in the case where the cleaner uptake rate of glucose is fixed to 0. (A) Heatmap of the productivity of the consortium (*DH*^*^) as a function of glucose inflow *G*_*in*_ and dilution rate *D*. The boundaries of the domains of coexistence and of existence of the sole producer are in dashed red and solid green lines, respectively. (B) For *G*_*in*_ = 20 g L^−1^, productivity as a function of *D* for the consortium (filled circles), and for a producer growing in isolation (empty circles). The color code for coexistence (dashed red) or existence of the sole producer (solid green) is the same as in the main text. Vertical dashed lines indicate different productivity domains (see discussion in main text). (C) Same as B for the process yield ((*DH*^*^)*/*(*DG*_*in*_)).

## S9 Text Relation between biomass degradation constant and maintenance coefficient^*^

In the main text, we use the relation *k*_*deg*_ = *C*_*m*_*Y*_*g*_ to estimate the value of the biomass degradation constant via the maintenance coefficient *C*_*m*_ and the biomass yield coefficient of glucose *Y*_*g*_. The degradation constant *k*_*deg*_ [h^−1^] expresses the non-growth-related maintenance rate per unit biomass, the maintenance coefficient *C*_*m*_ [g gDW^−1^ h^−1^] represents the rate of substrate taken up per unit biomass at zero growth rate and the yield coefficient *Y*_*g*_ [gDW g^−1^] characterizes the conversion of substrate (glucose) into biomass. Experimentally, the maintenance coefficient *C*_*m*_ is obtained by extrapolating the experimental data via the relation introduced by Pirt [1].

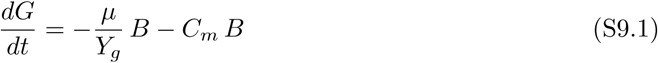

that relates the measured glucose uptake rate and the observed specific growth rate

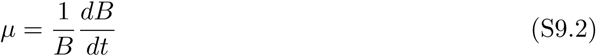

during exponential growth on glucose in batch. According to the relations introduced in the main text, in our model in batch conditions and in the presence of glucose as the sole carbon source, Eqs S9.1 and S9.2 simplify to

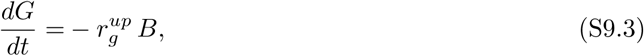

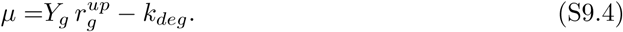

After substituting 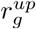 from Eq. S9.4 into Eq. S9.3, we obtain

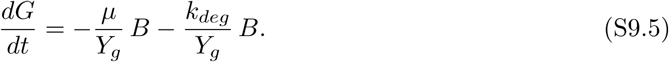

Compared to Eq. S9.1 this leads to the relation

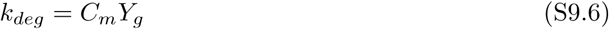

between the measured maintenance coefficient and the biomass degradation constant, used for computing the latter in the main text. Fig. S9.1 represents graphically Eqs S9.1 and S9.5. It is important to note that both *C*_*m*_ and *Y* might depend on the carbon source, while *k*_*deg*_ is independent of the substrate since it corresponds to the (negative) growth rate in the absence of nutrients. We denote the dependence of *C*_*m*_ and *Y* on glucose and acetate by the subscripts _*g*_ and _*a*_, respectively. Therefore, for a bacterial culture growing on acetate, the degradation rate

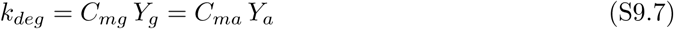

has the same value for a culture growing on glucose and acetate. In the case of acetate, one can produce a figure similar to Fig. S9.1. The slopes and the intercepts of the fitted line will be different for acetate, but the extrapolated lines must both go through the point (−*k*_*deg*_, 0).

**Figure S9.1:**
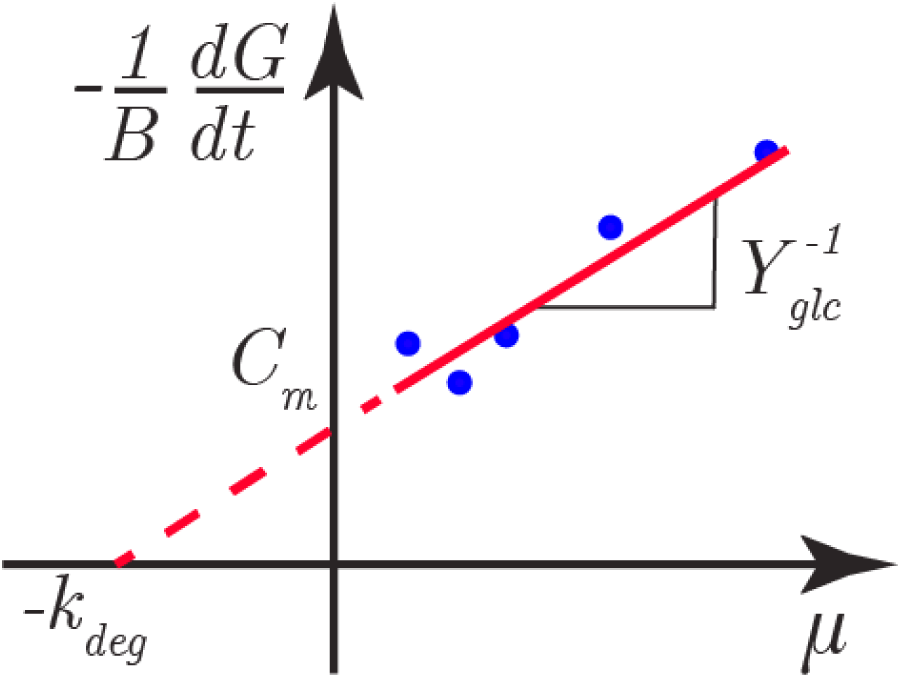
Relation between biomass degradation constant and maintenance coefficient. The maintenance coefficient *C*_*m*_ and the biomass degradation constant *k*_*deg*_ are obtained by extrapolating (dashed line) the data (dots) with Eqs S9.1 and S9.5, respectively, by measuring the specific substrate uptake rate −*dG/dt B*^−1^ as a function of the specific growth rate *µ* = *dB/dtB*^−1^. The yield coefficient *Yg* corresponds to the inverse of the slope of the interpolated relations (solid line).

## Footnotes

* Supporting Information of “Enhanced production of heterologous proteins by a synthetic microbial community: Conditions and trade-offs” (M. Mauri, J.-L. Gouzé, H. de Jong, E. Cinquemani)

